# Pharmacological hallmarks of allostery at the M4 muscarinic receptor elucidated through structure and dynamics

**DOI:** 10.1101/2022.09.27.509640

**Authors:** Ziva Vuckovic, Jinan Wang, Vi Pham, Jesse I. Mobbs, Matthew J. Belousoff, Apurba Bhattarai, Wessel A.C. Burger, Geoff Thompson, Mahmuda Yeasmin, Katie Leach, Emma T. van der Westhuizen, Elham Khajehali, Yi-Lynn Liang, Alisa Glukhova, Denise Wootten, Craig W. Lindsley, Andrew B. Tobin, Patrick M. Sexton, Radostin Danev, Celine Valant, Yinglong Miao, Arthur Christopoulos, David M. Thal

## Abstract

Allosteric modulation of G protein-coupled receptors (GPCRs) is a major paradigm in drug discovery. Despite decades of research, a molecular level understanding of the general principals that govern the myriad pharmacological effects exerted by GPCR allosteric modulators remains limited. The M_4_ muscarinic acetylcholine receptor (M_4_ mAChR) is a well-validated and clinically relevant allosteric drug target for several major psychiatric and cognitive disorders. Here, we present high-resolution cryo-electron microscopy structures of the M_4_ mAChR bound to a cognate G_i1_ protein and the high affinity agonist, iperoxo, in the absence and presence of two different positive allosteric modulators, LY2033298 or VU0467154. We have also determined the structure of the M_4_ mAChR-G_i1_ complex bound to its endogenous agonist, acetylcholine (ACh). Structural comparisons, together with molecular dynamics, mutagenesis, and pharmacological validations, have provided in-depth insights into the role of structure and dynamics in orthosteric and allosteric ligand binding, global mechanisms of receptor activation, cooperativity, probe-dependence, and species variability; all key hallmarks underpinning contemporary GPCR drug discovery.

## Introduction

Over the past 40 years, there have been major advances to the analytical methods that allow for the quantitative determination of the pharmacological parameters that characterise G protein-coupled receptor (GPCR) signaling and allosteric modulation (Figure 1A,B). These analytical methods are based on the operational model of agonism (Black and Leff, 1983) and have been extended or modified to account for allosteric modulation (Leach et al., 2007), biased agonism (Kenakin, 2012), and even biased allosteric modulation (Slosky et al., 2021). Collectively, these models and subsequent key parameters (Figure 1B) are used to guide allosteric drug screening, selectivity, efficacy and ultimately, clinical utility, and provide the foundation for modern GPCR drug discovery (Wootten et al., 2013). Yet, a systematic understanding of how these pharmacological parameters relate to the molecular structure and dynamics of GPCRs remains elusive.

**Figure 1.**
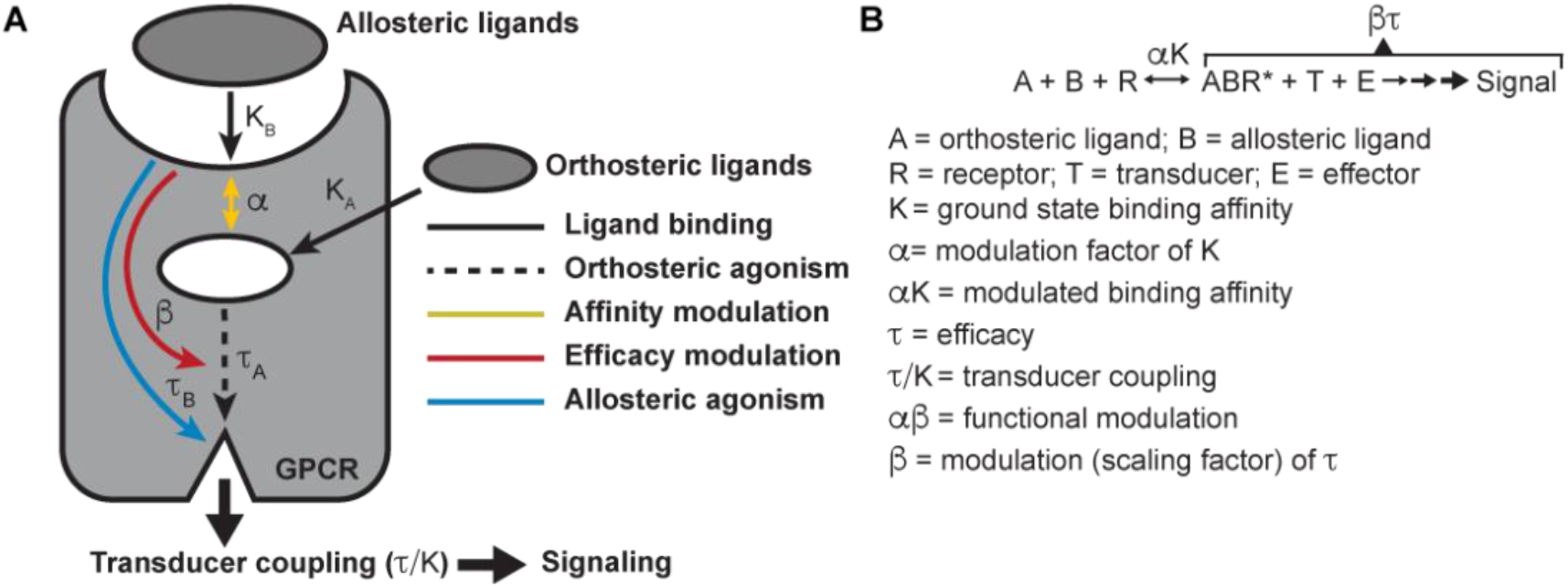
Pharmacological characterization of the PAMs, LY298 and VU154, with ACh and Ipx. (**A**) Schematic of the pharmacological parameters that define effects of orthosteric and allosteric ligands on a GPCR. (**B**) A simplified schematic diagram of the Black-Leff operational model to quantify agonism, allosteric, and agonist bias with pharmacological parameters defined (Black and Leff, 1983).

The muscarinic acetylcholine receptors (mAChRs) are an important family of five Class A GPCRs that have long served as model systems for understanding GPCR allostery (Conn et al., 2009). The mAChRs have been notoriously difficult to exploit therapeutically and selectively due to high sequence conservation within their orthosteric binding domains (Burger et al., 2018). However, the discovery of highly selective positive allosteric modulators (PAMs) for some mAChR subtypes has paved the way for novel approaches to exploit these high value drug targets (Chan et al., 2008; Gentry et al., 2014; Marlo et al., 2009). X-ray crystallography and cryo-electron microscopy (cryo-EM) have been used to determine inactive state structures for all five mAChR subtypes (Haga et al., 2012; Kruse et al., 2012; Thal et al., 2016; Vuckovic et al., 2019) and active state structures of the M_1_ and M_2_ mAChRs (Maeda et al., 2019). For the M_2_ mAChR this includes structures co-bound with the high-affinity agonist iperoxo (Ipx) and the PAM LY2119620 in complex with a G protein mimetic nanobody (Kruse et al., 2013) and the transducers G_o_ (Maeda et al., 2019) and β-arrestin1 (Staus et al., 2020). These M_2_ mAChR structures were foundational to validating the canonical mAChR allosteric site but are limited to only one agonist (iperoxo) and one PAM (LY2119620) and do not account for the vast pharmacological properties of ligands targeting mAChRs. A recent nuclear magnetic resonance (NMR) study at the M_2_ mAChR revealed differences in the conformational landscape of the M_2_ mAChR when bound to different agonists, but no clear link was established between the properties of the ligands and the conformational states of the receptor (Xu et al., 2019).

The M_4_ mAChR subtype is of major therapeutic interest due to its expression in regions of the brain that are rich in dopamine and dopamine receptors, where it regulates dopaminergic neurons involved in cognition, psychosis, and addiction (Bymaster et al., 2003; Dencker et al., 2011; Foster et al., 2016; Tzavara et al., 2004). Importantly, these findings have been supported by studies utilizing novel PAMs that are highly selective for the M_4_ mAChR (Bubser et al., 2014; Chan et al., 2008; Leach et al., 2010; Suratman et al., 2011). Among these, LY2033298 (LY298) was the first reported highly selective PAM of the M_4_ mAChR and displayed antipsychotic efficacy in a preclinical animal model of schizophrenia (Chan et al., 2008). Despite LY298 being one of the best characterized M_4_ mAChR PAMs, its therapeutic potential has been limited by numerous factors including its chemical scaffold, which has been difficult to optimize with respect to its molecular allosteric parameters (Figure 1) and variability of response between species (Suratman et al., 2011; Wood et al., 2017a). In the search for better chemical scaffolds, the PAM, VU0467154 (VU154), was subsequently discovered. VU154 showed robust efficacy in preclinical rodent models, however, it also exhibited species selectivity that prevented its clinical translation (Bubser et al., 2014). Collectively, LY298 and VU154 are exemplar tool molecules that highlight the promises and the challenges in understanding and optimising allosteric GPCR drug activity for translational and clinical applications.

Herein, by examining the pharmacology of the PAMs LY298 and VU154 with the agonists ACh and Ipx across radioligand binding assays and two different signaling assays and analysing these results with modern analytical methods, we determined the key parameters that describe signaling and allostery for these ligands. To investigate a structural basis for these pharmacological parameters, we used cryo-electron microscopy (cryo-EM) to determine high-resolution structures of the M_4_ mAChR in complex with a cognate G_i1_ heterotrimer and ACh and Ipx. We also determined structures of receptor complexes with Ipx co-bound with the PAMs LY298 or VU154. Moreover, because protein allostery is a dynamic process (Changeux and Christopoulos, 2016), we performed all-atom simulations using the Gaussian accelerated molecular dynamics (GaMD) enhanced sampling method (Draper-Joyce et al., 2021; Miao et al., 2015; Wang et al., 2021a) on the M_4_ mAChR using the cryo-EM structures. The structures and GaMD simulations, in combination with detailed molecular pharmacology and receptor mutagenesis experiments, provide fundamental insights into the molecular mechanisms underpinning the hallmarks of GPCR allostery. To further validate these findings, we investigated the differences in the selectivity of VU154 between the human and mouse receptors and established a structural basis for species selectivity. Collectively, these results will enable future GPCR drug discovery research and potentially lead to the development of next generation M_4_ mAChR PAMs.

## Results

### Pharmacological characterisation of M_4_ mAChR PAMs with ACh and Ipx

*Hallmarks of ligand binding.* We first used radioligand binding assays (Figure 2A) to determine the *ground state binding affinities* of ACh and Ipx (K_A_) for the orthosteric site and of LY298 and VU154 (K_B_) for the allosteric site of the unoccupied human M_4_ mAChR (Figure 2B), along with the degree of *binding cooperativity* (α) between the agonists and PAMs when the two are co-bound (Figure 2C). Analysis of these experiments revealed that LY298 and VU154 have very similar binding affinities for the allosteric site with values (expressed as negative logarithms; pK_B_) of 5.65 ± 0.07 and 5.83 ± 0.12, respectively (Figure 2B), in accordance with previous studies (Bubser et al., 2014; Leach et al., 2011). Both PAMs potentiated the ground state binding affinity of ACh and Ipx (Figure 2D,E), with the effect being greatest between LY298 and ACh (approx. 400-fold increase in binding affinity). Comparatively, the positive cooperativity between VU154 and ACh was only 40-fold. When Ipx was used as the agonist, the binding affinity modulation mediated by both PAMs was more modest, characterized by an approximate 72-fold potentiation for the combination of Ipx and LY298, and 10-fold potentiation for the combination of Ipx and VU154. These results indicate *probe-dependent* effects (Valant et al., 2012) with respect to the ability of either PAM to modulate the affinity of each agonist (Figure 2D,E). It is important to note that binding modulation is *reciprocal* and the affinities of LY298 and VU154 were also increased in the agonist bound state (Figure 2E). This results in LY298 having a 5-fold higher binding affinity than VU154 when agonists are bound (Table S1).

**Figure 2.**
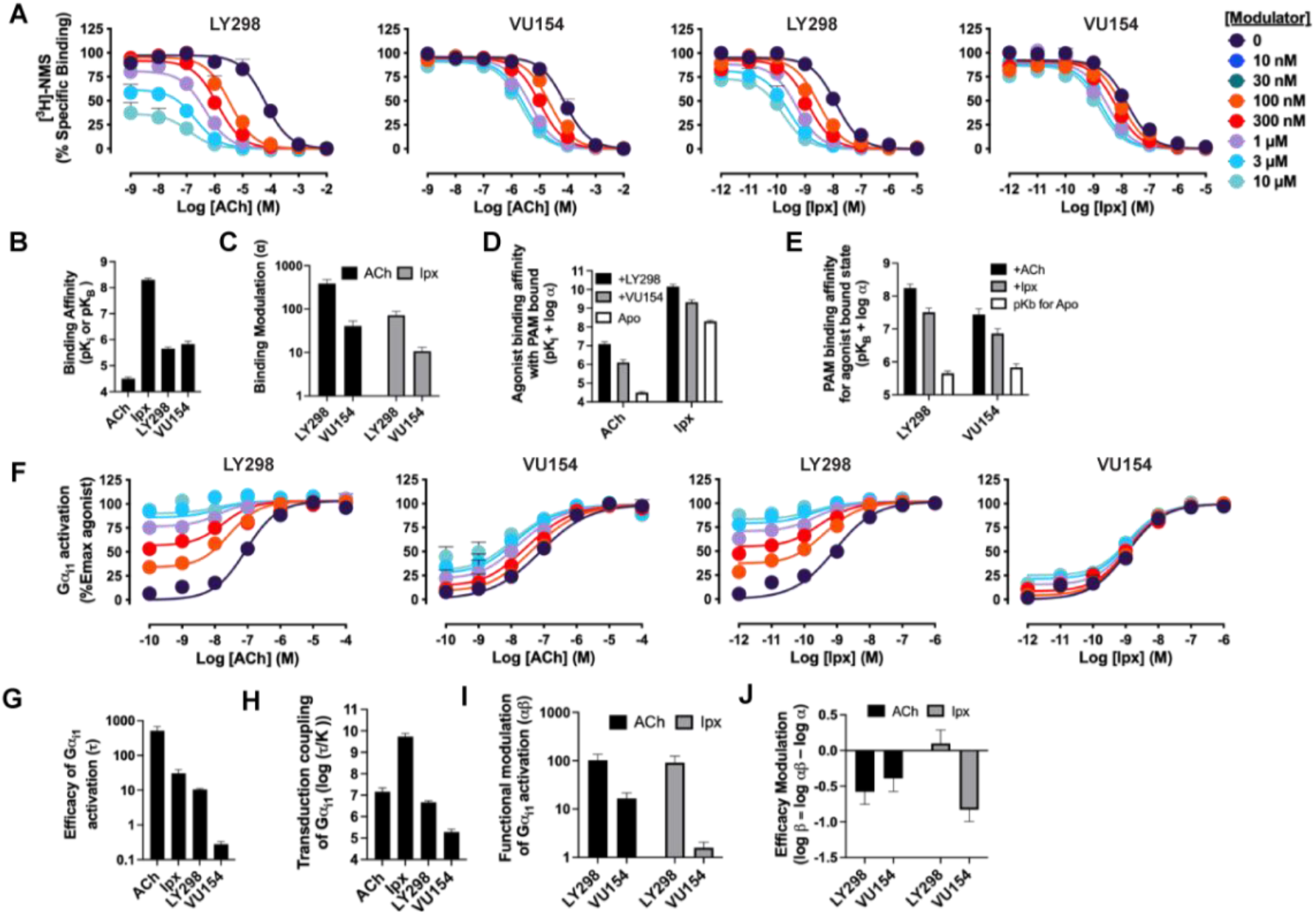
Pharmacological characterization of the PAMs, LY298 and VU154, with ACh and Ipx. (**A**) Concentration response curves of interactions between the orthosteric and allosteric ligands at the human M_4_ mAChR in [^3^H]-N-methylscopolamine ([^3^H]-NMS) binding assays.(**B-E**) Quantification of data from (**A**) to calculate (**B**) equilibrium binding affinities (pK_i_ and pK_B_), (**C**) the degree of binding modulation (α) between the agonists and PAMs, and the modified affinities (**D**) αK_A_ and (**E**) αK_B_. (**F**) Concentration response curves of interactions between the orthosteric and allosteric ligands at the human M_4_ mAChR with an area under the curve analysis of Gα_i1_ activation using the TruPath assay. (**G-J**) Quantification of data from (**A,F**) to calculate (**G**) the signaling efficacy (τ_A_ and τ_B_) and (**H**) the the transduction coupling coefficients (log (τ/K)) of each ligand, (**I**) the functional cooperativity (αβ) between ligands, and (**J**) the efficacy modulation (β) between ligands. All data are mean ± SEM of 3 or more independent experiments performed in duplicate or triplicate with the pharmacological parameters determined from a global fit of the data. The error in (**D,E,J**) was propagated using the square root of the sum of the squares. See Table S1.

#### Hallmarks of GPCR function

We subsequently used the BRET-based TruPath assay (Olsen et al., 2020), as a proximal measure of G protein activation with Gα_i1_ (Figure 2F). We also used a more amplified downstream signalling assay, extracellular signal-regulated kinases 1/2 phosphorylation (pERK1/2), that is also dependent on G_i_ activation (Figure S1A), to measure the cell-based activity of each PAM with each agonist. These signaling assays allowed us to determine the *efficacy* of the agonists (τ_A_) and the PAMs (τ_B_) (Figure 2G, Figure S1B). Importantly, efficacy (τ), as defined from the Black-Leff operational model of agonism (Black and Leff, 1983), is determined by receptor density (B_max_), the ability of an agonist to promote an active receptor conformation, and the ability of a cellular system to generate a response (Figure 1B). Notably, in both signalling assays, the rank order of efficacy was ACh > Ipx > LY298 > VU154. We subsequently calculated the *transducer coupling coefficient* (τ/K) (Figure 1B; Figure 2H; S1C), a parameter often used to quantify agonist bias. The transducer coupling coefficient accounts for the ground state binding affinity of the agonist (K), either orthosteric or allosteric, and characterises the agonism of a specific pathway defined as the interaction between an agonist, receptor, and transducer, which indirectly interacts with effector and signaling proteins (Kenakin et al., 2012). Accordingly, in both assays, the rank order of transducer coupling was Ipx >> ACh ∼ LY298 > VU154 due to Ipx having a higher ground state binding affinity for the receptor. Overall, these results indicate that although ACh is a more efficacious agonist than Ipx it has lower transducer coupling coefficient. In contrast, LY298 has both better efficacy and transducer coupling than VU154 (Table S1).

The signaling assays and use of an operational model of allosterism also allowed for the determination of the *functional cooperativity* (αβ) exerted by the PAMs (Figure 2I; S1D), which is a composite parameter accounting for both binding (α) and efficacy (β) modulation. Notably, VU154 displayed lower positive functional cooperativity with ACh than LY298. Strikingly, VU154 had negligible functional modulation with Ipx in contrast to the cooperativity observed with ACh in the TruPath assay. The 10-fold difference in αβ values for VU154 between ACh and Ipx highlights the dependence of the orthosteric probe used in the assay (i.e. *probe dependence*); on this basis, VU154 would be classified as a NAL (not a PAM) with Ipx in the TruPath assay (Table S1).

The degree of *efficacy modulation* (β) that the PAMs have on the agonists can be calculated directly by subtracting the binding modulation (α) from the functional modulation (αβ) (Figure 2J; S1E). A caveat of this analysis is that errors for β are higher due the error being propagated between experiments. Ideally, the degree of efficacy modulation would be determined in an experimental system where the maximal efficacy of system is not reached by the agonists alone (Berizzi et al., 2016). Nevertheless, our analysis shows the PAMs LY298 and VU154 appear to have a slight negative to neutral effect on agonist efficacy in the G_i1_ Trupath and pERK1/2 assays (Table S1), suggesting that the predominant allosteric effect exerted by these PAMs is mediated through binding modulation.

Collectively, our extensive analysis on the pharmacology of LY298 and VU154 with ACh and Ipx offers detailed insight into the key differences between these ligands across a range of pharmacological properties: ligand binding, probe dependence, efficacy, agonist-receptor-transducer interactions, and allosteric modulation (Figure 1, Table S1). We hypothesised that structures of the human M_4_ mAChR in complex with different agonists and PAMs combined with molecular dynamic simulations could provide high resolution molecular insights into the different pharmacological profiles of these ligands.

### Determination of M_4_R-G_i1_ complex structures

Similar to the approach used in prior determination of active-state structures of the M_1_ and M_2_ mAChRs (Maeda et al., 2019), we used a human M_4_ mAChR construct that lacked residues 242 to 387 of the third intracellular loop to improve receptor expression and purification, and made complexes of the receptor with G_i1_ protein and either the endogenous agonist, ACh, or Ipx. Due to the higher affinity of Ipx compared to ACh (Schrage et al., 2013), we utilised Ipx to form additional M_4_R-G_i1_ complexes with or without the co-addition of either LY298 or VU154. In all instances, complex formation was initiated by combining purified M_4_ mAChR immobilized on anti-FLAG resin with detergent solubilized G_i1_ membranes, a single-chain variable fragment (scFv16) that binds G_i_ and Gβ, and the addition of apyrase to remove guanosine 5’-diphosphate (Maeda et al., 2018). For this study, we used a G_i1_ heterotrimer composed of a dominant negative form of human Gα_i1_, and human Gβ_1_ and Gγ_2._ (Liang et al., 2018a). Vitrified samples of each complex were imaged using conventional cryo-TEM on a Titan Krios microscope (Danev et al., 2021).

The structures of ACh-, Ipx-, LY298-Ipx-, and VU154-Ipx-bound M_4_R-G_i1_ complexes were determined to resolutions of 2.8, 2.8, 2.4, and 2.5 Å, respectively (Figure 3A-E, S2, Table S2). For the ACh-bound M_4_R-G_i1_ complex, an additional focus refinement yielded an improved map of the receptor and binding site (2.75 Å) for modelling (Figure 3E). The electron microscopy (EM) density maps for all complexes were sufficient for confident placement of backbone and sidechains for most of the receptor, G_i1_, and scFv16, and the bound ligands with exception of the alkyne bond of Ipx (Figure S3). Notably, in the Ipx-bound structures, EM density for the alkyne bond of Ipx was missing (Figure S3F), matching previous cryo-EM Ipx-bound mAChR structures (Maeda et al., 2019). As such, it is difficult to place the alkyne bond of Ipx into one preferred pose, largely because of rotational freedom on the carbon between the alkyne bond and the rotatable trimethyl ammonium ion. This is highlighted by the different poses of the alkyne bond across the different Ipx-bound mAChR structures and is consistent with the reported docking of Ipx in the M_2_ mAChR structure (Figure S3F) (Maeda et al., 2019).

**Figure 3.**
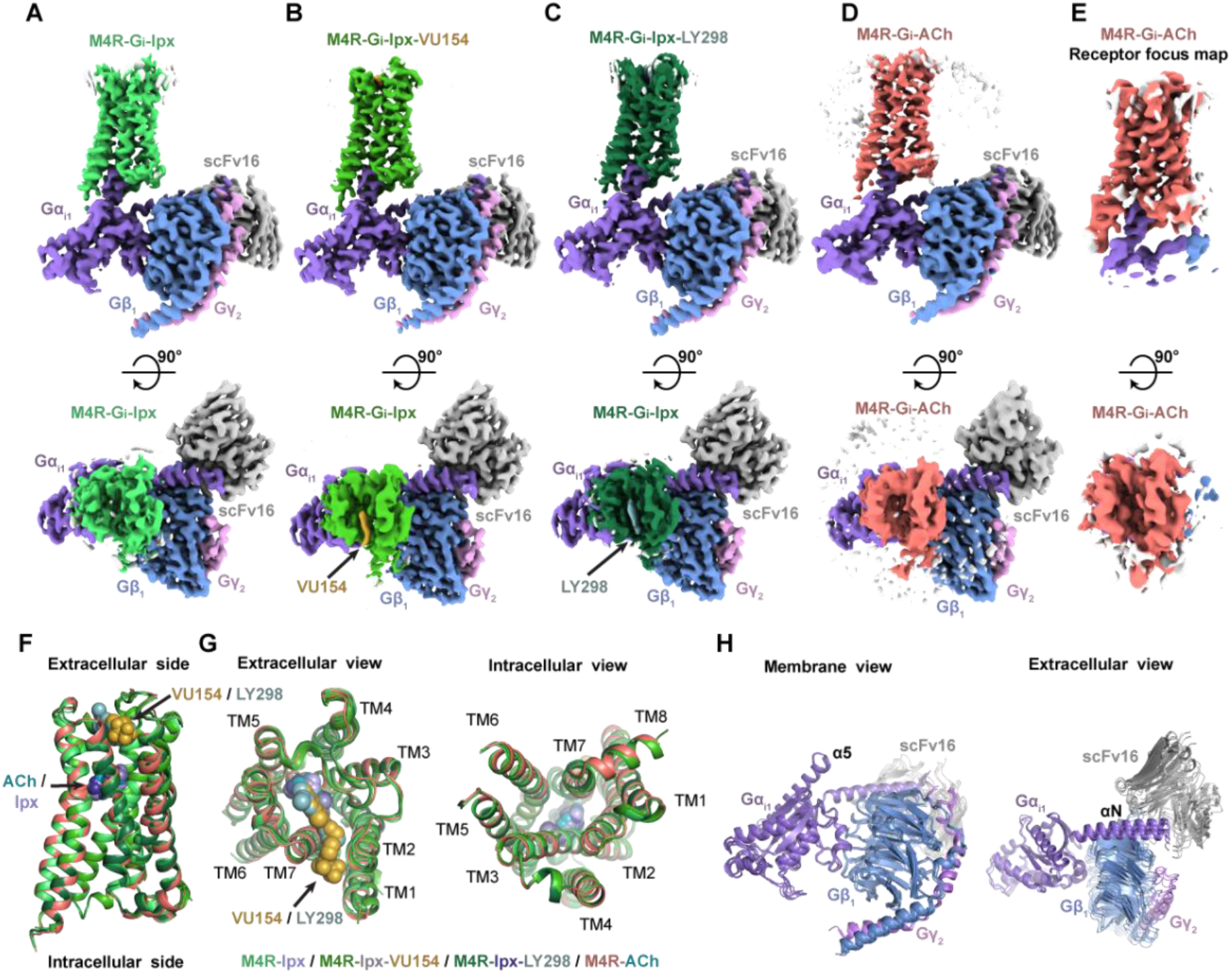
Cryo-EM structures of the M_4_R-G_i1_-scFv16 complexes. (**A-E**) Cryo-EM maps of (**A**) Ipx-bound, (**B**) VU154-Ipx-bound, (**C**) LY298-Ipx bound, and (**D,E**) ACh-bound M_4_R-G_i1_-scFv16 complexes with views from the membrane and the extracellular surface. (**F,G**) Comparison of the receptor models with bound ligands and views from the (**G**) extracellular and intracellular surface of the receptors. (**H**) Comparison of the positions of Gα_i1_Gβ_1_Gγ_2_-scFv16 with views from the membrane and extracellular surface.

In all four structures, EM density beyond the top of transmembrane helix 1 (TM1) and the third intracellular loop (ICL3) of the receptor was poorly observed and not modelled. Similarly, the EM density of the α-helical domain of Gα_i1_ was poor and not modelled. These regions are highly dynamic and typically not modelled in many class A GPCR-G protein complex structures. Apart from these regions, most amino acid side chains were well resolved in the final EM density maps (Figure S3).

### Structure and dynamics of agonist binding

Recently, cryo-EM structures of M_4_R-G_i1_ complexes bound to ipx, ipx and the PAM LY2119620, and an allosteric agonist c110, were determined (Wang et al., 2022). Surprisingly, comparison of the M_4_R-G_i1_ complex structures reveal larger differences in the position of key orthosteric and allosteric site residues than the M_1_R-G_11_ and M_2_R-G_oA_ complex structures (Figure S4-5). Unfortunately, the quality of density in the EM maps around the orthosteric and allosteric sites of these M_4_R-G_i1_ structures (Wang et al., 2022) was poor resulting in several key residues being mismodelled in each site (Figure S5). Therefore, differences between the M_4_R-G_i1_ structures are highly likely to not be due to genuine differences, and as such we compared to the M_1_R-G_11_ and M_2_R-G_oA_ complex structures in this study (Maeda et al., 2019).

Overall, our M_4_R-G_i1_ complex structures are similar in architecture to that of other activated class A GPCRs including the M_1_R-G_11_ and M_2_R-G_oA_ complexes (Figure S4). Superposition of the M_4_R-G_i1_ complexes revealed nearly identical structures with root mean square deviations (RMSD) of 0.4–0.5 Å for the full complexes and 0.3–0.4 Å for the receptors alone (Figure 3F). The largest differences occur around the extracellular surface of the receptors (Figure 3G) along with slight displacements in the position of the αN helix of Gα_i1_ and Gβ_1_, Gγ_2_, and scFv16 with respect to the receptor (Figure 3H). The EM density of side chains surrounding the ACh and Ipx binding sites (Figure 4A-B) was well resolved providing the opportunity to understand structural determinants of orthosteric agonist binding. The orthosteric site of the M_4_ mAChR, in common with the other mAChR subtypes, is buried within the TM bundle in an aromatic cage that is composed of four tyrosine residues, two tryptophan residues, one phenylalanine residue, and seven other polar and nonpolar residues (Figure 4C). Notably, all 14 of these residues are absolutely conserved across all five mAChR subtypes, underscoring the difficulty in developing highly subtype-selective orthosteric agonists (Burger et al., 2018). Both ACh and Ipx have a positively charged trimethyl ammonium ion that makes cation-π interactions with Y113^3.33^, Y416^6.51^, Y439^7.39^, and Y443^7.43^ (Figure 4C) (superscript refers to the Ballesteros and Weinstein scheme for conserved class A GPCR residues) (Ballesteros and Weinstein, 1995). Likewise, both ACh and Ipx have a polar oxygen atom that can form a hydrogen bond to the indole nitrogen of W164^4.57^ with the oxygen of Ipx also being in position to interact with the backbone of N117^3.37^ (Figure 4D). Mutation of any of these contact residues reduces the affinity of ACh, validating their importance for agonist binding (Leach et al., 2011; Thal et al., 2016). The largest chemical difference between ACh and Ipx is the bulkier heterocyclic isoazoline group of Ipx that makes a π-π interaction with the conserved residue W413^6.48^ (Figure 4D). The residue W413^6.48^ is part of the CWxP motif, also known as the rotamer toggle switch, a residue that typically undergoes a change in rotamer between the inactive and active states of class A GPCRs (Shi et al., 2002).

**Figure 4.**
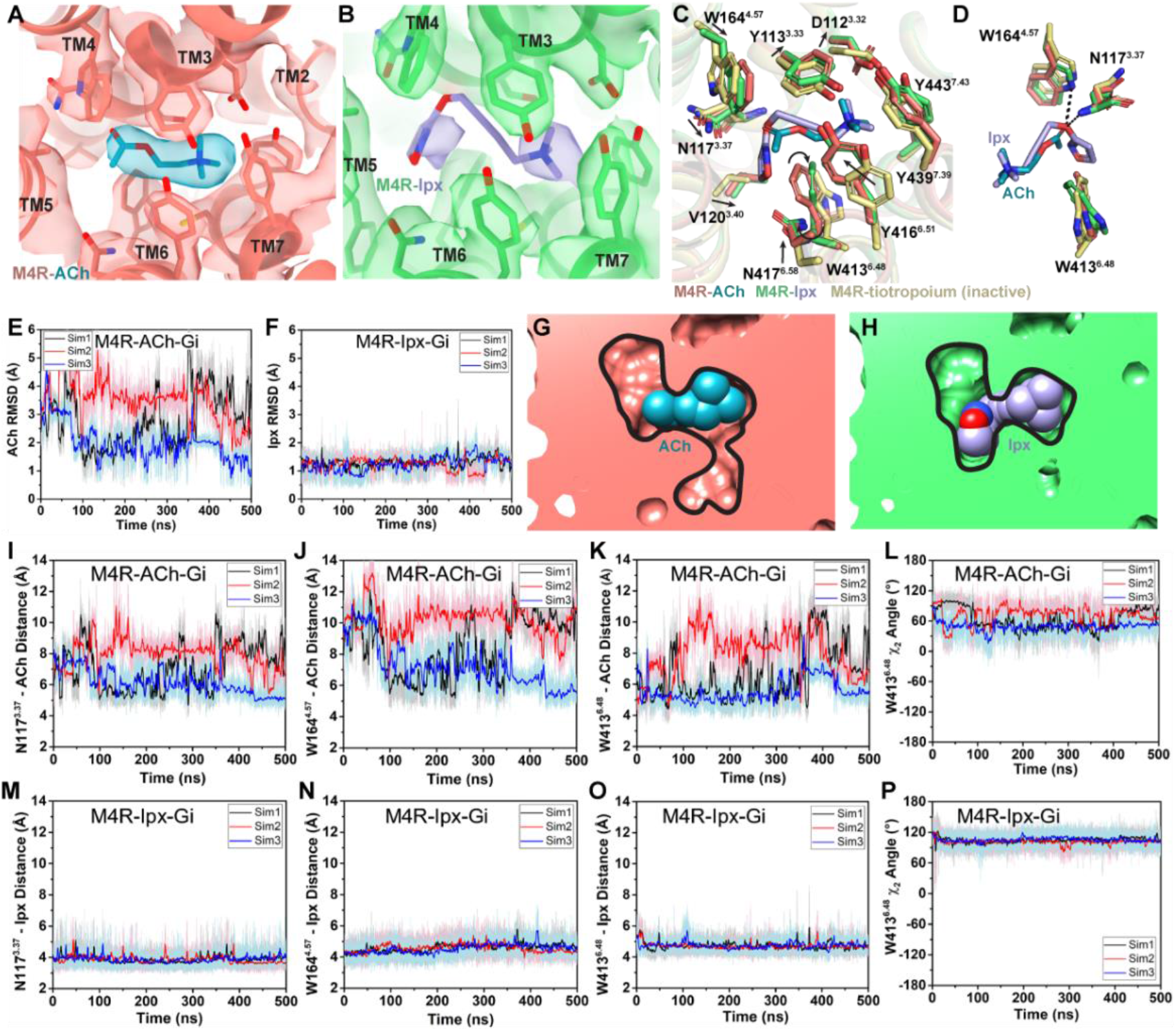
Interactions of ACh and Ipx with the receptor. (**A,B**) Cryo-EM density of the (A) ACh- and (**B**) Ipx-bound structures. (**C,D**) Interactions at the orthosteric binding site comparing the active state ACh- and Ipx-bound structures with the inactive state tiotropium bound structure (PDB: 5DSG). Arrows denote relative movement of residues between the inactive and active states. (**D**) Detailed interactions of ACh and Ipx. Hydrogen bonds are shown as black dashed lines. (**E-F, I-P**) Time courses from GaMD simulations of the ACh- and Ipx-bound M_4_R-G_i1_ cryo-EM structures, each performed with 3 separate replicates. Individual replicate simulations are illustrated with different colours. The heading of each plot refers to the specific model used in the simulations. RMSDs of (**E**) ACh and (**F**) Ipx from simulations of the cryo-EM structures. (**G,H**) Cross-sections through the ACh- and Ipx-bound structures denoting the relative size of the binding pockets outlined in black. (**I-P**) The distances of interactions between ACh and Ipx with residues (**I, M**) N117^3.37^, (**J,N**) W164^4.67^, and (**K,O**) W413^6.48^, and (**L,P**) the χ2 angle of W413^6.48^. See Table S3.

To investigate the structural dynamics of the M_4_ mAChR, we performed three independent 500 ns GaMD simulations on the ACh- and Ipx-bound M_4_R-G_i1_ cryo-EM structures (Table S3). GaMD simulations revealed that ACh undergoes higher fluctuations in the orthosteric site than Ipx (Figure 4E,F). Similarly, the interactions of N117^3.37^, W164^4.57^, and W413^6.48^ with Ipx were more stable than those with ACh (Figure 4I-P). In the ACh-bound structure, W413^6.48^ was in a conformation that more closely resembled the inactive-state tiotropium-bound structure (Figure 4C,D). GaMD simulations also showed that W413^6.48^ sampled a larger conformational space in the ACh-bound structure than in the Ipx-bound structure (Figure 4L,P). The predominate χ_2_ angle of W413^6.48^ was approximately 60^◦^ and 105^◦^ in the ACh-bound and Ipx-bound simulations, respectively, corresponding to the cryo-EM conformations.

Located above ACh and Ipx is a tyrosine lid formed by three residues (Y113^3.33^, Y416^6.51^, and Y439^7.39^) that separates the orthosteric binding-site from an extracellular vestibule (ECV) at the top of the receptor and the bulk solvent (Figure 4C). In the inactive conformation, the tyrosine lid is partially open due to Y416^6.51^ rotating away from the binding pocket to accommodate the binding of bulkier inverse agonists such as tiotropium. In contrast, mAChR agonists are typically smaller in size than antagonists and inverse agonists, and this is reflected in a contraction of the size of the orthosteric binding pocket from 115 Å^3^ when bound to tiotropium to 77 and 63 Å^3^ when bound to ACh and Ipx, respectively (Figure 4G,H) (Tian et al., 2018). Together, the smaller binding pocket of Ipx and more stable binding interactions with nearby residues that include W413^6.48^ likely explain why Ipx has greater than 1,000-fold higher binding affinity than ACh.

### Structure and dynamics of PAM binding and allosteric modulation of agonist affinity

The M_4_R-G_i1_ structures of LY298 and VU154 co-bound with Ipx are very similar to the Ipx- and ACh-bound structures, as well as to prior structures of the M_2_ mAChR bound to Ipx and the PAM, LY2119620 (Figure S4) (Kruse et al., 2013; Maeda et al., 2019). Both LY298 and VU154 bind directly above the orthosteric site in the ECV that is composed of a floor delineated by the tyrosine lid, and ‘walls’ formed by residues from TM2, TM6, TM7, ECL2, and ECL3 (Figure 5A,B). The EM density surrounding the PAM binding site and the ECV of the M_4_ mAChR were clearly resolved with one exception; in the VU154-bound structure the EM density begins to weaken around the trifluoromethylsulfonyl moiety (Figure 5A). This was likely due to the moiety’s ability to freely rotate (similarly to the alkyne bond of Ipx in the orthosteric site) and a lack of strong interactions with the receptor.

**Figure 5.**
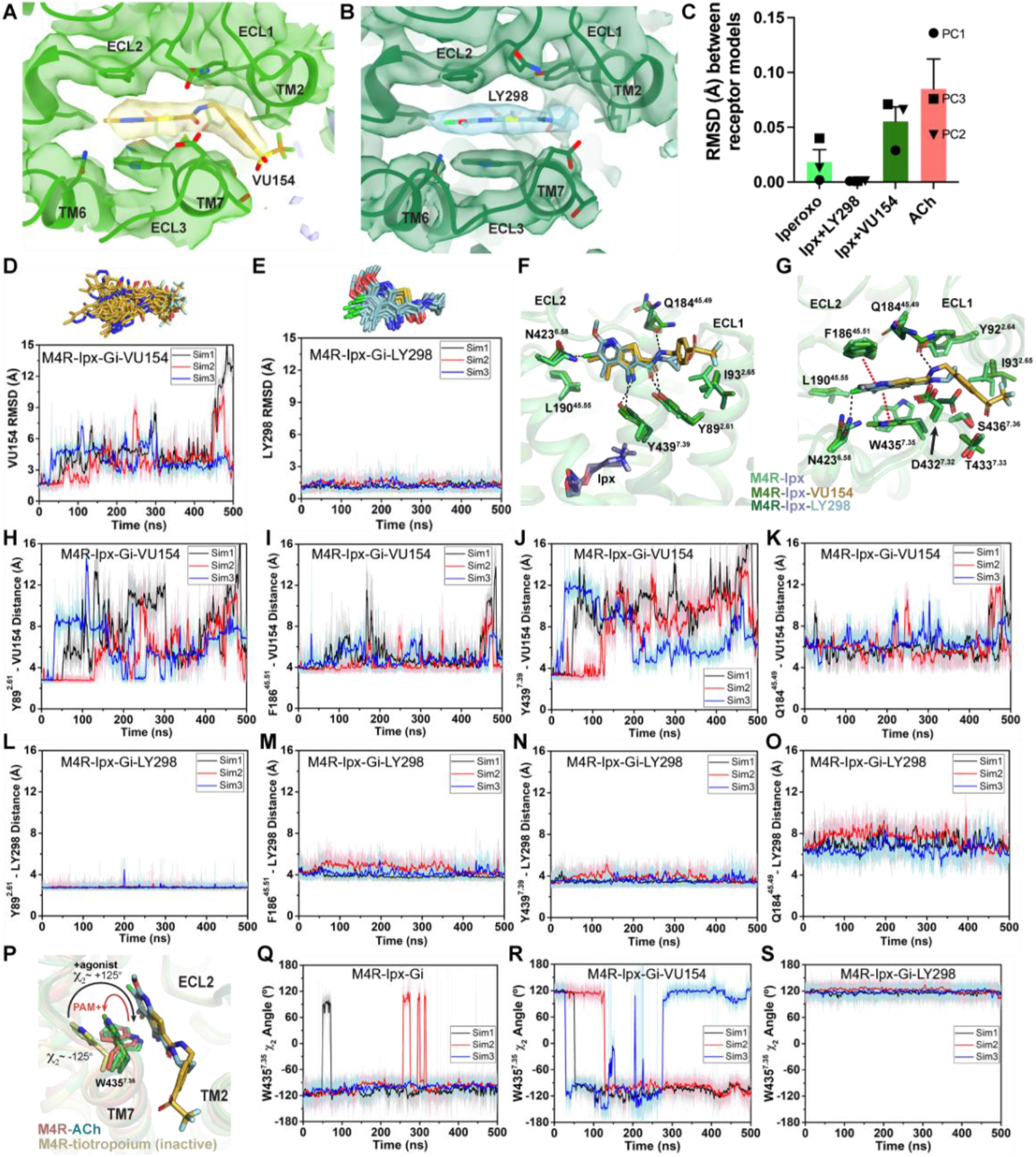
Binding and dynamics of LY293 and VU154. (**A,B**) Cryo-EM density of the (**A**) VU154- and (**B**) LY298-binding sites. (**C**) The root-mean square deviations (RMSD) between receptor models of the respective cryo-EM structures that were refined into the first and last frames of the EM maps from each principal component (PC1-PC3) of the 3D variability analysis. Values shown are mean ± SEM. (**D,E**) Top representative binding conformations of (**D**) VU154 aI(**E**) LY298 obtained from structural clustering with frame populations ≥ 1% and time courses of the RMSDs of each PAM relative to the cryo-EM structures. (**F,G**) Binding interactions of VU154 and LY298 with views from the (**F**) membrane and (**G**) extracellular surface. (**H-O**) Time-courses from three 500 ns GaMD simulations using the (**H-K**) VU154- and (**L-O**) LY298-Ipx-bound cryo-EM structures. Distances between the interactions of VU154 and LY298 with residues (**H, L**) Y89^7.39^, (**I,M**) F186^45.51^, Y439^7.39^, and Q184^45.49^. (**P**) Position and χ2 angle of W435^7.35^ in the tiotropium-, ACh-, Ipx-, VU154-Ipx-, and LY298-Ipx bound structures. (**Q-S**) Time courses of the W435^7.35^ χ2 angle obtained from GaMD simulations in the (**Q**) I-, (**R**) VU154-Ipx-, and (S) LY298-Ipx-bound cryo-EM structures. See Table S3.

Given the overall similarities revealed by our four cryo-EM structures, we examined if there were further differences in the dynamics between the PAM-bound structures by performing a 3D multivariance analysis (3DVA) of the principal components of motion within the Ipx-, LY298-Ipx, VU154-Ipx, and ACh-bound M_4_R-G_i1_ cryo-EM data sets using Cryosparc (Punjani and Fleet, 2021); a similar analysis performed previously on cryo-EM structures of class A and class B GPCRs provided important insights into the allosteric motions of extracellular domains and receptor interactions with G proteins (Josephs et al., 2021; Liang et al., 2020; Mobbs et al., 2021; Zhang et al., 2020).

In the 3DVA of the Ipx-bound complex, the M4 receptor appeared less flexible than the receptor in the ACh-bound complex (**Supplemental Videos 1-2**) consistent with Ipx having a higher binding affinity and more stable pose during the GaMD simulations (Figure 4E,F). The LY298-Ipx-bound complex appeared similar to the Ipx-bound complex with LY298 being bound in the ECV (**Supplemental Video 3**). In contrast, the 3DVA of the VU154 structure had more dynamic movements in the allosteric pocket that could reflect partial binding of VU154 (**Supplemental Video 4**). This observation was in line with our findings that VU154 had lower binding cooperativity (Figure 2C) and functional cooperativity with agonists than LY298 (Figure 2I; S1D). To quantify the differences from the 3DVA, we rigid body fitted and refined the respective M_4_R-G_i1_ models into the first and last frames of the EM maps from each principal component of the 3DVA and then calculated the RMSD between the receptor models from the from the first and last frames (Figure 5C). In agreement with our prior observations, the VU154-Ipx-bound and ACh-bound complexes had greater RMSDs with values of 0.06 and 0.09 Å respectively. Comparatively, the Ipx-bound and LY298-Ipx-bound complexes had lower RMSD values of 0.02 and 0.001 Å, respectively. The results of the 3DVA do not represent *bona fide* measures of receptor dynamics, rather they are suggestive of there being differences between the collected data sets that led to the structures. To support these findings, we compared the GaMD simulations of all four cryo-EM structures (Table S3). Notably, VU154 underwent considerably higher fluctuations than LY298 with RMSDs ranging from 1.5–15 Å for VU154 and 0.8–2.1 Å for LY298 relative to the cryo-EM structures (Figure 5D,E). Therefore, the GaMD simulations corroborate our 3DVA results and suggests that complexes bound to agonists with high affinity or co-bound with agonists and PAMs with high positive cooperativity will exhibit lower dynamic fluctuations.

To investigate why the binding of LY298 was more stable than VU154, we examined the ligand interactions with the receptor. There are three key binding interactions that are shared between both PAMs and the M_4_ mAChR: 1) a three-way π-stacking interaction between F186^45.51^ (ECL2 residues have been numbered 45.X denoting their position between TM4 and TM5 with X.50 being a conserved cysteine residue), the aromatic core of the PAMs, and W435^7.35^; 2) a hydrogen bond between Y439^7.39^ of the tyrosine lid and the primary amine of the PAMs; and 3) a hydrogen bond between Y89^2.61^ and the carbonyl oxygen of the PAMs (Figure 5F,G). While these interactions are conserved for both PAMs in the consensus cryo-EM maps, during GaMD simulations these interactions were more stable with LY298 than VU154 (Figure 5H-O). The importance of these interactions was validated pharmacologically (Figure S6; Table S4), whereby mutation of any of these residues completely abolished the binding affinity modulation mediated by LY298 and VU154 at the M_4_ mAChR with both Ipx and ACh as agonists.

A potential fourth interaction was observed with residue Q184^45.49^ and the amide nitrogen of the PAMs; however, the GaMD simulations suggest that this interaction is relatively weak (Figure 5K,O), consistent with the fact that mutation of Q184^45.49^ to alanine had no effect on the binding modulation of LY298 or VU154 (Figure S6; Table S4). In addition, each PAM has at least one unique binding interaction with the receptor (Figure 5F,G). For LY298, this is an interaction between the fluorine atom and N423^6.58^ that appeared to be stable during simulation and, when mutated to alanine reduced the binding modulation of LY298 (Figure S7A) (Thal et al., 2016). For VU154, two additional hydrogen bonding interactions were formed with Y92^2.64^ and T433^7.33^ (Figure 5G), although these interactions were fluctuating during GaMD simulations (Figure S7B,C). Finally, W435^7.35^ is a key residue in the ECV that changes from a planar rotamer in the agonist-bound structures to a vertical rotamer that π stacks against the PAMs (Figure 5P). In GaMD simulations of the Ipx-bound structure, W435^7.35^ is predominantly in a planar conformation that corresponds to its conformation in the cryo-EM structure (Figure 5P,Q). In contrast, the binding of LY298 stabilizes W435^7.35^ into a vertical position (Figure 5P,S). However, in the VU154-bound receptor, W435^7.35^ appears to alternate between the planar and vertical positions, consistent with VU154 having a less stable binding pose (Figure 5R). These results indicate that the binding of LY298 is more stable than VU154 due to LY298 being able to form stable binding interactions with key residues in the ECV. This provides a likely explanation for why LY298 was able to exert greater positive binding cooperativity on orthosteric agonists than VU154.

### A molecular mechanism of probe dependence

As highlighted above, PAMs, LY298 and VU154, displayed stronger allosteric binding modulation with ACh than Ipx, an example of probe dependence (Figure 2, S1D). These findings are in accord with previous studies where we identified probe dependence in the actions of LY298 when tested against other orthosteric agonists (Chan et al., 2008; Suratman et al., 2011). To investigate a mechanism for probe dependence at the M_4_ mAChR, we performed GaMD simulations with LY298 and VU154 co-bound with ACh, by replacing Ipx with ACh in the corresponding cryo-EM structures (Table S3 and Figure S7). In the absence of PAM, ACh was more dynamic than Ipx with root-mean-square fluctuations (RMSF) of 2.13 Å versus 0.88 Å, reflective of the fact Ipx binds with higher affinity than ACh (Figure S7L). In the presence of LY298 or VU154 the dynamics of ACh binding was decreased, with RMSFs reduced to 1.23 Å and 1.82 Å, respectively, and with LY298 having the greatest effect (Figure S7L). This is in line with LY298 having more cooperativity with ACh than VU154 (Figure 2C). In comparison to ACh, there was a modest increase in the dynamics of Ipx with the addition of LY298 or VU154, likely reflecting the fact Ipx binding to the receptor was already stable (Figure S7D-F). These results provide a plausible mechanism for probe dependence, at least with regards to differences in the magnitude of the allosteric effect depending on the ligand bound. Namely, PAMs manifest higher cooperativity when interacting with agonists, such as ACh, that are inherently less stable on their own when bound to the receptor, in contrast to more stable ligands such as Ipx.

### Structural and dynamic insights into orthosteric and allosteric agonism

In addition to the ability to allosterically modulate the function of orthosteric ligands, it has become increasingly appreciated that allosteric ligands may display variable degrees of direct agonism in their own right, over and above any allosteric modulatory effects (Changeux and Christopoulos, 2016). Prior studies have established that the activation process of GPCRs involves conformational changes that extend from the extracellular domains through to the intracellular surface (Nygaard et al., 2009). Comparison of the active state ACh-, Ipx-, LY298-Ipx-, and VU154-Ipx-bound M_4_R-G_i1_ structures to the inactive state tiotropium-bound M_4_ mAChR structure (Protein Data Bank accession 5DSG) (Thal et al., 2016) thus affords an opportunity to gain new insights into the activation process mediated by multiple orthosteric agonists in the presence and absence of two different PAMs that display high (LY298) and low (VU154) degrees of direct allosteric agonism (Figure 1F, 6A-C).

**Figure 6.**
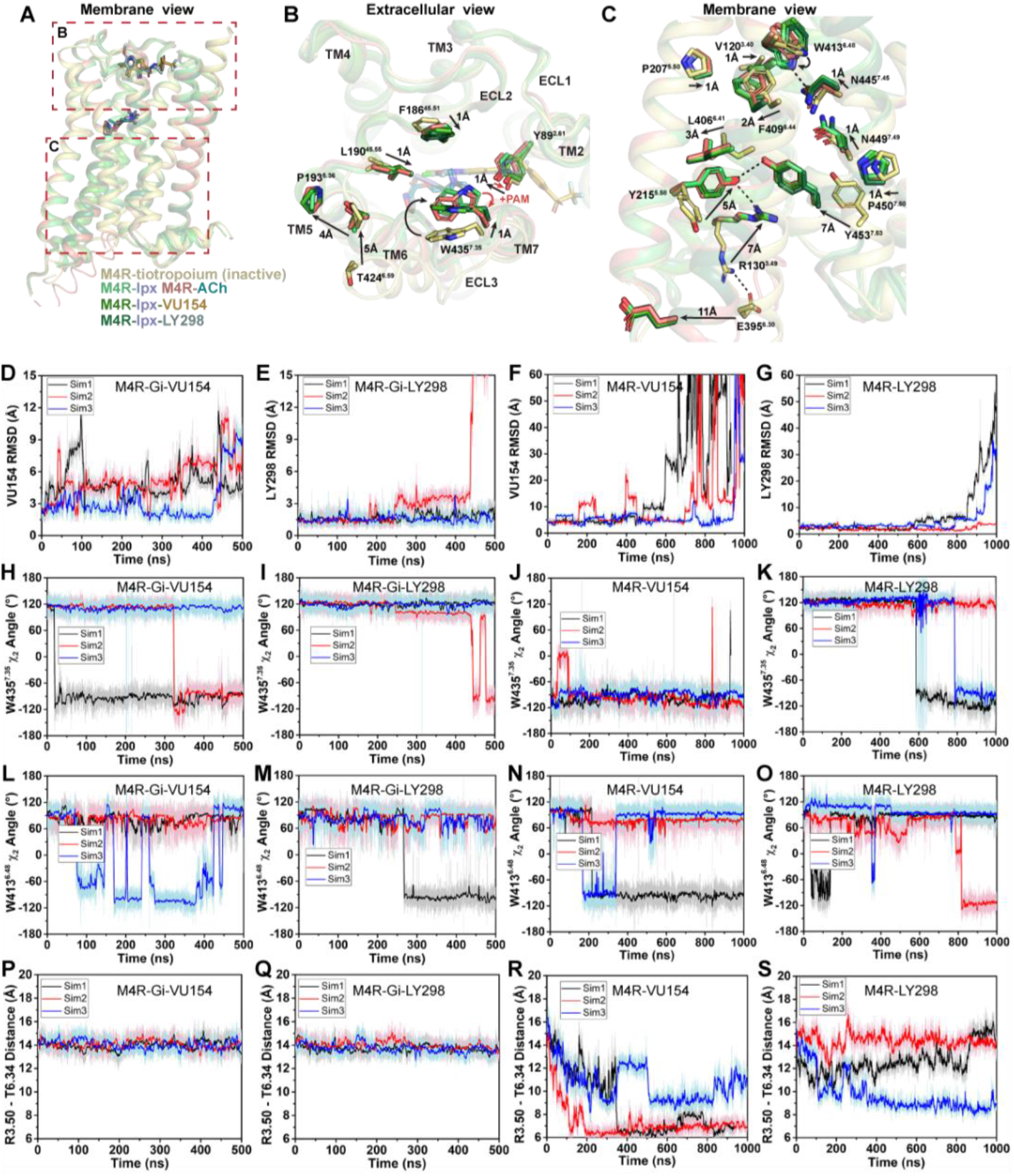
Structural and dynamic insights into orthosteric and allosteric agonism. (**A**) Cartoon of the receptor models indicating regions of interest for panels (**B-C)** shown within the red boxes. (**B**) View of the tiotropium-bound, agonist-bound, and PAM-agonist-bound conformations from the extracellular surface. (**C**) Membrane view of residues and activation motifs involved in signalling. (**D-S**) Time courses of (**D-G**) the PAMs RMSDs, (**H-K**) the W435^7.35^ χ2 angle, (**L-O**) the W413^6.48^ χ2 angle, and (**P-S**) the TM3-TM6 distance measured by distance between R130^3.50^ and T399^6.34^ obtained from GaMD simulations of the M_4_R-G_i1_-VU154, M_4_R-G_i1_-LY298, M_4_R-VU154, and M_4_R-LY298 systems, respectively. See Table S3.

As previously discussed, agonist binding decreases the size of the orthosteric binding site (Figure 4G,H). The primary driver of this decrease was the tyrosine lid residue Y416^6.51^, which underwent a large rotation towards Y113^3.33^ creating a hydrogen bond that seals off the tyrosine lid (Figure 4C). The closure of the tyrosine lid was further reinforced by a change in the rotamer of W435^7.35^ to a planar position that sits parallel to the tyrosine lid allowing for a π-π interaction with Y416^6.51^ and a positioning of the indole nitrogen of W435^7.35^ to potentially form a hydrogen bond with the hydroxyl of Y89^2.61^ (Figure 6B). The contraction of the orthosteric pocket by the inward movement of Y416^6.51^ also led to a contraction of the ECV with a 5 Å inward movement of the top of TM6 and ECL3. As a consequence, the top of TM5 was displaced outward by 4 Å forming a new interface between TM5 and TM6 that was stabilized by a hydrogen bond between T424^6.59^ and the backbone nitrogen of P193^5.36^ along with aromatic interactions between F197^5.40^ and F425^6.60^ (Figure 6B). These interactions were specific to the active state structures and appear to be conserved as they were also present in the M_1_ and M_2_ mAChR active state structures (Maeda et al., 2019). In addition to the movements of TM5 and TM6, there was a smaller 1 Å inward movement of ECL2 (Figure 6B). The binding of LY298 and VU154 had minimal impact on the conformation of most ECL residues, implying that the reorganisation of residues in the ECV by orthosteric agonists contributes to the increased affinity of the PAMs (Figure 2E). There was a slight further inward shift of ECL2 towards the PAMs to facilitate the 3-way π-stacking interaction with F186^45.51^ and W435^7.35^. In addition, in the PAM-bound structures, Y89^2.61^ rotated away from its position in the ACh-and Ipx-bound structures either due to a loss of an interaction with W435^7.35^ or to form a better hydrogen bond with the carbonyl oxygen of the PAMs (Figure 6B).

Below the orthosteric binding site are several signaling motifs that are important for the activation of class A GPCRs, including the PIF motif (Rasmussen et al., 2011; Wacker et al., 2013), the Na^+^ binding site (Liu et al., 2012b; White et al., 2018), the NPxxY motif (Fritze et al., 2003), and the DRY motif (Figure 6C) (Ballesteros et al., 2001). The conformations of these activation motifs were very similar across all four active-state M_4_ mAChR structures and were consistent with the position of these motifs across other active-state class A GPCR structures (Zhou et al., 2019). Collectively, all of the described activation motifs facilitate an 11 Å outward movement of TM6 that typifies GPCR activation and creation of the G protein binding site. In comparison to the ECV residues (Figure 6B), beyond the rotamer toggle switch residue W413^6.48^, there are no discernible differences between the agonist and PAM-agonist-bound structures, suggesting a shared activation mechanism for residues below W413^6.48^ (Figure 6C).

As indicated above, LY298 also displays robust allosteric agonism in comparison to VU154 (Figure 2G,H). To probe whether the allosteric agonism of LY298 could be related to its ability to better stabilize the M_4_ mAChR in an active conformation in comparison to VU154, we performed additional GaMD simulations on the LY298-Ipx- and VU154-Ipx-bound M_4_R-G_i1_ structures with the agonist Ipx removed (3x 500 ns) and with both Ipx and the G protein removed (3x 1,000 ns) (Figure 6, Table S3). In GaMD simulations, LY298 underwent lower RMSD fluctuations than VU154 before dissociating from the receptor (Figure 6D-G). Similarly, the conformations of W435^7.35^ and W413^6.48^ were better stabilized in the LY298-Ipx-bound systems, indicating that LY298 more strongly promotes an active receptor conformation (Figure 6H-K). In the presence of the G protein, both PAMs stabilised an active conformation of the receptor based on the distances between TM3 and TM6 (Figure 6P,Q). Upon removal of the G protein, the VU154-bound M_4_ mAChR quickly transitioned towards the inactive conformation, while the LY298-bound M_4_ mAChR was more resistant to deactivation in the GaMD simulations (Figure 6R,S). This observation supports LY298 having greater efficacy than VU154 (Figure 2G) as it better stabilizes the active conformation of the M_4_ mAChR. Overall, the GaMD simulations show that in the absence of agonist alone, or agonist and G protein, LY298 better stabilizes activation motifs from the top of the receptor (W435^7.35^) all the way down to the intracellular G protein binding pocket (DRY-TM6) providing mechanistic insights into the function of LY298 as a stronger PAM-agonist than VU154.

### A molecular basis of species selectivity

One of the main advantages of allosteric modulators is the ability to selectivity target highly conserved proteins. The mAChRs are prime example where allosteric modulators have been designed to selectivity target specific subtypes. To date the only PAM-bound mAChR structures are ones with LY2119620, a PAM that has activity at both the M_2_ and M_4_ mAChRs. Similarly, LY298 has activity at the M_2_ mAChR. However, the allosteric properties of VU154 are differentially affected by the species of the receptor (Wood et al., 2017a, 2017b). At the human M_4_ mAChR, LY298 displays robust binding modulation, functional modulation, and allosteric agonism, while VU154 has comparatively weaker allosteric properties (Figure 1, Table S1). Conversely, at the mouse M_4_ mAChR, VU154 has a high degree of positive binding modulation, functional modulation, and allosteric agonism that is comparable to LY298 at the human M_4_ mAChR (Figure S7, Table S1). Therefore, we aimed to determine if our prior findings could be used to explain the selectivity of VU154 between the human and mouse receptors.

The amino acid sequences of the human and mouse M_4_ mAChRs are highly conserved, with most of the differences occurring between the long third intracellular loop and the N- and C-termini. As shown in Figure 7A, only three residues differ between the human and mouse M_4_ mAChR with respect to the transmembrane domain. Specifically, residue V91 (L in mouse) at the top of TM2 points into the lipid bilayer, and D432 and T433 (E and R in mouse), which are located at the top of TM7 and form part of the allosteric binding site near VU154.

**Figure 7.**
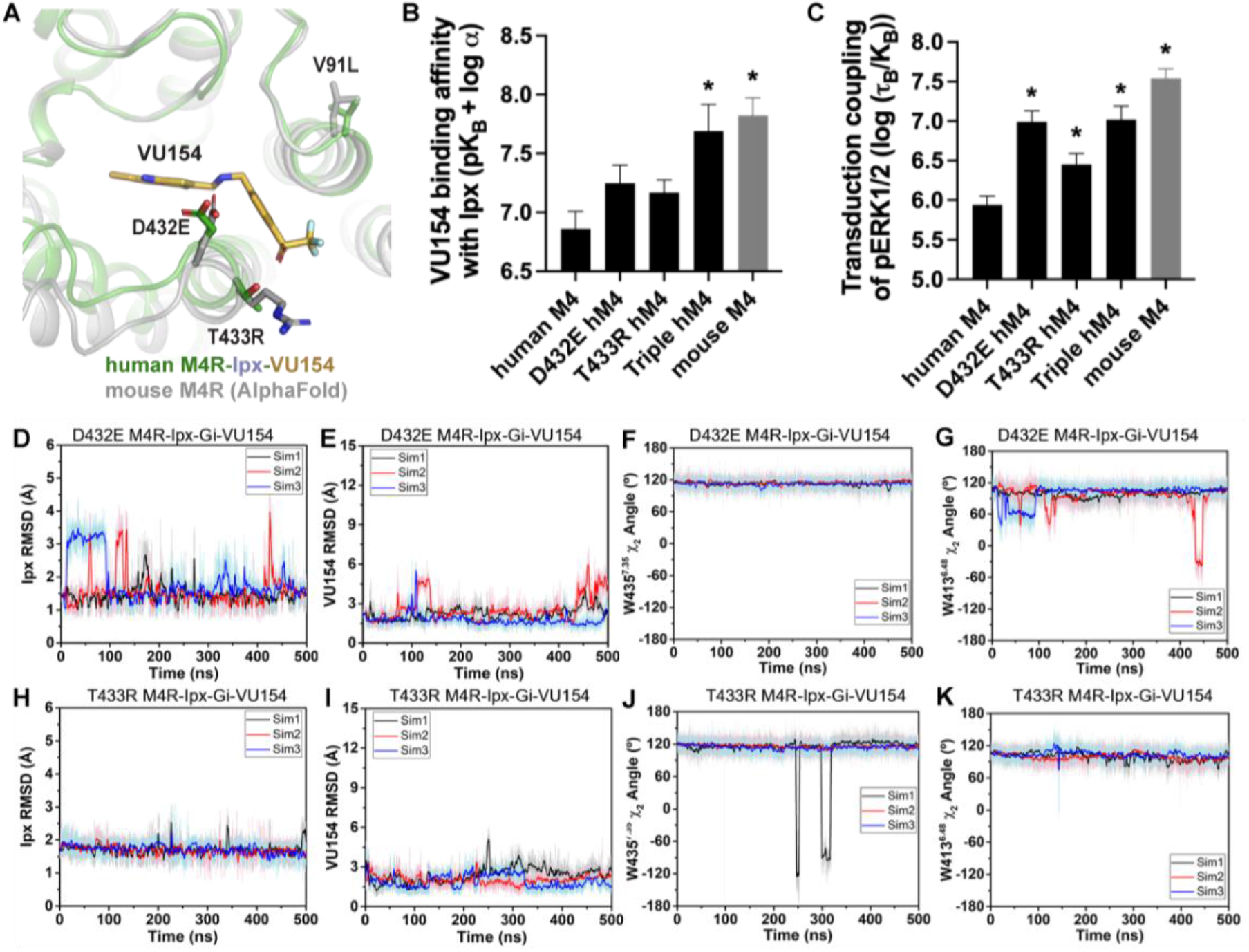
A molecular mechanism for the species selectivity for VU154. (**A**) Comparison of the cryo-EM structure of the human M4 mAChR bound to Ipx-VU154 with the AlphaFold model of the mouse M4 mAChR (Jumper et al., 2021; Varadi et al., 2022). The three residues that differ between species and within the core 7TM bundle from the human receptor (V91, D432, and T433) are shown as sticks along with the corresponding residues from the mouse receptor. (**B**) The binding affinity of VU154 for the Ipx-bound conformation (pK_B-Ipx_ = pK_B_ + α) determined from [^3^H]-NMS binding experiments. Values calculated with data from **Figure S8A** with propagated error. (**C)** Transduction coupling coefficients (log (τ/K)) of pERK1/2 signaling from data from **Figure S8A,E**). (**D-K**) Time courses of obtained from GaMD simulations of the (**D-G**) D432E and (**H-K**) T433R mutant M_4_R-Ipx-G_i1_-VU154 systems with (**D,H**) Ipx RMSDs, (**E,I**), VU154 RMSDs, (**F,J**) W435^7.35^ χ_2_ angle, and (**G,K**) W413^6.48^ χ_2_ angle. See Table S4 and Figure S8. (**B,C**) Data shown are mean ± SEM from 3 or more experiments performed in duplicate with the pharmacological parameters determined from a global fit of the data. *Indicates statistical significance (p < 0.05) relative to WT as determined by a one-way ANOVA with a Dunnett’s post-hoc test.

Previous work suggested that residues D432 and T433 were important for differences in the species selectivity of LY298 (Chan et al., 2008). As such, we examined two single D432E and T433R mutants and a V91L/D432E/T433R triple mutant of the human receptor, along with the mouse M_4_ mAChR in radioligand binding and pERK1/2 experiments using Ipx and both PAMs (Figure S8, Table S1). For LY298, there were no statistically significant differences in binding or function between species and across the mutants that were more than 3-fold in effect. In contrast, VU154 had a 10-fold higher binding affinity for the Ipx-bound mouse M_4_ mAChR (compare Figure 2EG with Figure 7B). The affinity of VU154 increased by 2.5-fold at the D432E and T433R mutants and the triple mutant matched the affinity of the mouse receptor (Figure 7B). In functional assays, similar results were observed for VU154 with Ipx at the mouse M_4_ mAChR, with significant increases in the efficacy (τ_B_ – corrected for receptor expression), transduction coefficients (τ_B_/K_B_), and the functional modulation (αβ) (Figure 7B**, S8,** Table S1). Relative to the WT M_4_ mAChR, the efficacy, transduction coefficients, and functional modulation of VU154 increased for all of the mutants (Figure S8), however, none of the values fully matched the mouse receptor. Nevertheless, these results indicate that V91L, D432E, and T433R play a key role in mediating the species selectivity of VU154.

Our prior findings suggest the robust allosteric activity of LY298 at the human M_4_ mAChR was due to stable interactions with the receptor (Figure 5). As a proof-of-principle, we questioned if GaMD simulations would produce a stable binding mode for VU154 with D432E and T433R mutations to the VU154-Ipx-bound M_4_R-G_i1_ cryo-EM structure that was similar to our previously observed stable binding pose of LY298 (Figure 5, Table S3). Excitingly, both the D432E and T433R mutants resulted in a dynamic profile of VU154 that matched our GaMD simulations of LY298 from the LY298-Ipx-bound M_4_R-G_i1_ cryo-EM structure, including stabilized VU154 binding, constrained χ_2_ rotamer conformations of W435^7.35^ and W413^6.48^, and stable binding interactions with Y89^2.61^, Y439^7.39^, Q184^45.49^, and F186^45.51^ (Figure 7D-K, S8, **Supplemental Video 9,10**). Collectively, these findings reiterate the importance of receptor dynamics in the determination of allosteric modulator selectivity, as even subtle differences in amino acid residues between species may result in profound changes in overall stability of the same PAM-agonist-receptor complex.

## Discussion

Major advances have been made in recent years in the appreciation of the role of GPCR allostery and its relevance to modern drug discovery (Changeux and Christopoulos, 2016; Wootten et al., 2013). Despite an increase in the number of reported high-resolution GPCR structures bound to allosteric ligands (Thal et al., 2018), there remains a paucity of molecular-level details about the interplay between the complex chemical and pharmacological parameters that define allostery at GPCRs. By combining detailed pharmacology studies, multiple high-resolution cryo-EM structures of the M_4_ mAChR bound to two distinctly pharmacologically different agonists and PAMs, and GaMD simulations, we have now provided exquisite in-depth insights into the relationship between both structure and dynamics that govern multiple facets of GPCR allostery (Figure 8A).

**Figure 8.**
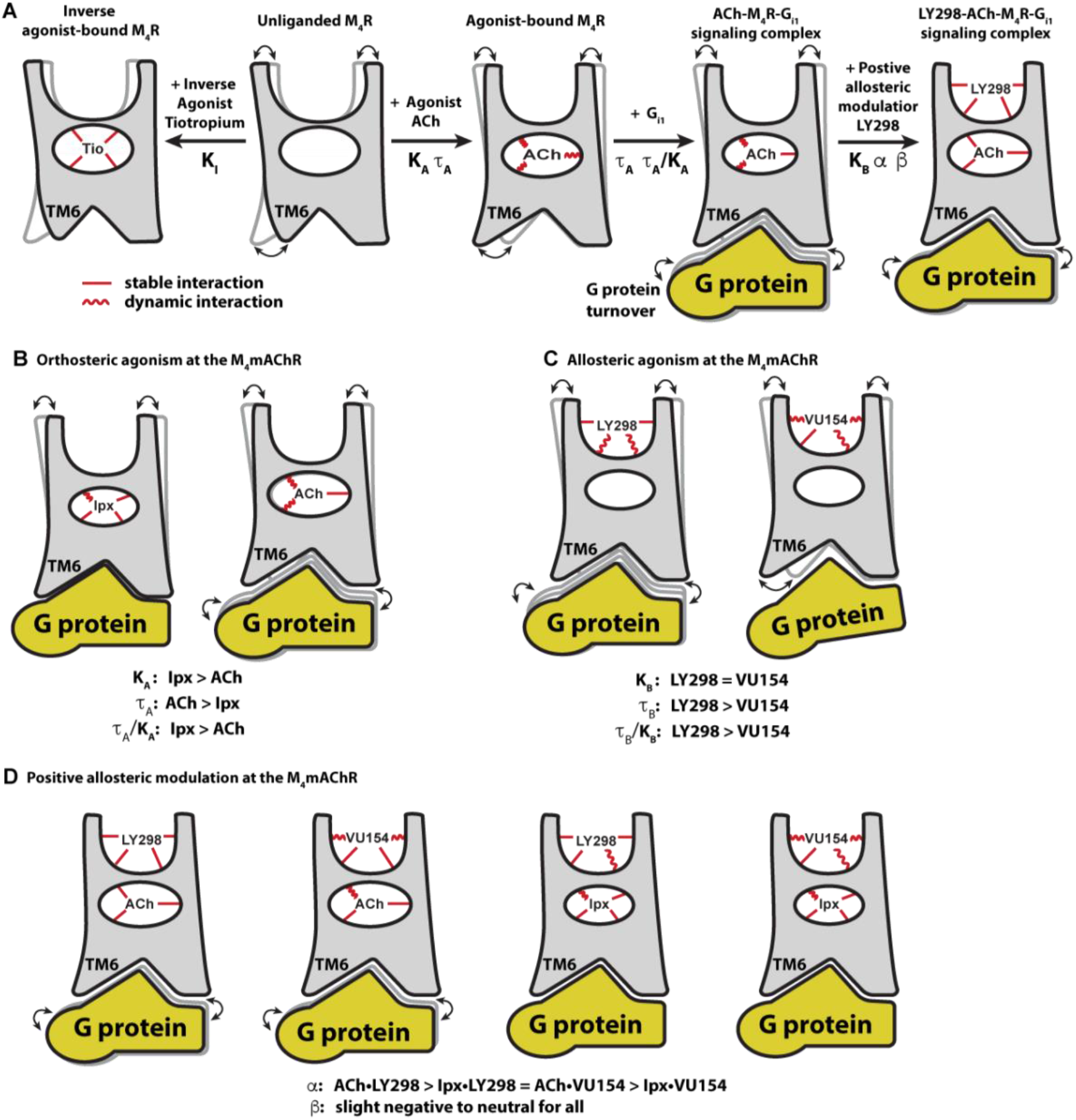
Conformational dynamics of the allostery at M_4_ mAChR signalling complexes. (**A**) A schematic cartoon illustrating the conformational states of the ligands and the M_4_ mAChR when bound to different types of ligands and transducer, along with the resulting dynamic profiles. Pharmacological parameters related to each conformational change are shown. Stable ligand-receptor interactions are denoted by a straight line and less-stable (more dynamic) interactions are denoted by a wavy line. (**B**) Ipx bound the M_4_ mAChR with a higher affinity and more stability than ACh but had lower efficacy. ACh being more loosely bound and coupled to G protein may facilitate more G protein turnover accounting for its higher efficacy. (**C**) LY298 and VU154 bound to the M_4_ mAChR with similar affinity for the ground of the receptor, but LY298 was found to bind more stably. LY298 had a higher efficacy than VU154, suggesting that allosteric agonism at the M_4_ mAChR is mediated by stabilization of the ECV.(**D**) The PAMs LY298 and VU154 display robust binding modulation at the M_4_ mAChR with LY298 having a stronger allosteric effect. Both PAMs displayed stronger binding modulation with the agonist ACh versus Ipx, an example of probe dependence. Both PAMs also displayed a slight negative to neutral effect on the efficacy of the agonists, suggesting that their mechanism of action is largely through binding.

Comparison of the ACh- and Ipx-bound M_4_ mAChR structures revealed that Ipx bound in a smaller binding pocket (Figure 4G,H), and GaMD simulations showed that Ipx formed more stable interactions with the receptor (Figure 4I-P). These observations likely explained why Ipx exhibited greater than 1,000-fold higher binding affinity than ACh (Figure 2B), being consistent with studies of other agonists at the β_1_-adrenoceptor and the M_1_ mAChR (Brown et al., 2021; Warne et al., 2019) (Figure 8B). The observation that ACh was a more efficacious agonist than Ipx (Figure 2G) yet bound with lower affinity and less stable interactions than Ipx was paradoxical. Kenakin and Onaran previously opined on the paradox between ligand binding affinity and efficacy (Kenakin and Onaran, 2002), and showed via simulations that, in general, there was a negative correlation between binding affinity and efficacy. One interpretation of these results was that the ACh-bound M_4_ mAChR more readily sampled receptor conformations that engaged with the transducers (Manglik et al., 2015). Similarly, the ACh-bound M_4_ mAChR may also have faster G protein turnover than Ipx due to Ipx-M_4_R-Gi1 forming a more stable ternary complex (Furness et al., 2016) (Figure 8B). In fact, the later point was supported by Ipx having a greater transducer coupling coefficient than ACh (Figure 2H) and suggests that structures of GPCRs in a ternary complex (agonist-receptor-transducer) are better represented by their transducer coupling coefficients than the efficacy of the agonist with respect to the transducer mediated signalling pathway(s). Given that transducer coupling coefficients were used to calculate ligand bias, one might expect structures of GPCR ternary complexes to capture conformations associated with ligand bias. Indeed, structures of GPCRs bound to biased ligands have now been reported for many GPCRs (Liang et al., 2018b; Masureel et al., 2018; McCorvy et al., 2018; Wacker et al., 2013; Warne et al., 2012; Wingler et al., 2019). Currently, however, there is no generalizable structural basis that explains ligand bias (Seyedabadi et al., 2022) as the conformational differences are likely to be more subtle and dynamic, and therefore require the combination of techniques like NMR spectroscopy, single molecule FRET, and MD simulations to fully resolve (Cao et al., 2021; Cong et al., 2021; Gregorio et al., 2017; Huang et al., 2021; Katayama et al., 2021; Liu et al., 2012a; Solt et al., 2017; Sušac et al., 2018; Ye et al., 2016).

This study highlighted that two PAMs with distinctly different pharmacological profiles (Figure 1) may bind to and stabilize receptor conformations that were very similar when viewed as static structures (Figure 5). In contrast, the 3DVA analysis from our cryo-EM structures suggested differences in the dynamics of the cryo-EM structures that were explored further in GaMD simulations (Figure 5C**, Supplementary Videos 1-4**) and revealed that LY298 had a more stable binding pose and interactions with the receptor than VU154 in the PAM-agonist-receptor-transducer bound conformation. These observations were consistent with LY298 having greater positive binding cooperativity than VU154 (Figure 2E) and suggested that GaMD simulations of PAMs bound to the M_4_ mAChR could be an extremely valuable tool for future drug discovery and optimization (Bhattarai and Miao, 2018).

Pharmacological analysis revealed that LY298 is a better PAM-agonist than VU154 with respect to efficacy (Figure 2G) and transduction coefficients (Figure 2H) in the G_i1_ Trupath and pERK1/2 signalling assays. GaMD simulations of the PAM-receptor-transducer and PAM-receptor bound complexes, again showed that LY298 more stably interacted with the receptor (Figure 5) and in the absence of G protein better stabilized the duration of the active conformation of the receptor (Figure 6). These findings were not contradictory to our above findings that ACh was more efficacious than Ipx despite having weaker interactions with the receptor, because when the ground state affinity of the ligands was accounted for in the transduction coupling coefficients the rank order of agonism was Ipx >> ACh ∼ LY298 > VU154 (Figure 2B). Furthermore, these results were in accordance with the observations of Kenakin and Onaran that ligands with the same binding affinity can also have differing efficacies (and vice-versa). In addition, the mechanism of agonism for allosteric ligands that bind to the ECV may differ (Xu et al., 2021). Prior work by DeVree *et al*. established that allosteric coupling of G proteins to the unliganded active receptor conformation promoted closure of the ECV region (DeVree et al., 2016). This allosteric coupling is reciprocal and stabilizing the ECV region by PAMs likely leads to increased efficacy (Figure 8C).

The PAMs LY298 and VU154 also displayed stronger allosteric effects with ACh than with Ipx, an observation known as probe dependence (Figure 2C-E). Probe dependence can have substantial implications on how allosteric ligands are detected, validated, and their potential therapeutic utility (Kenakin, 2005). Examples of probe dependence are not limited to studies on mAChRs and have been observed across multiple receptor families (Christopoulos, 2014; Gentry et al., 2015; Pani et al., 2021; Slosky et al., 2020; Wang et al., 2021b). GaMD simulations comparing the PAMs co-bound to either Ipx or ACh showed that the PAMs had a stabilizing effect on ACh, whereas the stability of Ipx was slightly reduced by the PAMs likely because the binding of Ipx was already stable. This is a sensible explanation from thermodynamic principles. Another explanation invokes the two-state receptor model (Canals et al., 2011), which stipulates that the degree of positive modulation for PAMs increases with an increase in the efficacy of the agonists. The pharmacology data support this model, as ACh was more efficacious than Ipx and was better modulated by both PAMs (Figure 8D). These results again highlight the apparent paradox between ligands being more stably bound to the receptor but also having lower efficacy. Supporting both observations is the fact that the positive functional modulation (αβ) of LY298 and VU154 in G_i1_ signalling assays is driven by positive binding modulation (α), as the efficacy modulation (β) of the PAMs was negative to neutral (Figure 2J, S1E). These observations were consistent with recent studies that suggest the conformational dynamics between agonist and receptor were important for functional signalling (Bumbak et al., 2020; Cary et al., 2022; Deganutti et al., 2022; O’Connor et al., 2015).

The findings presented here provide new insight into the allosteric signalling and allosteric modulation of GPCRs by combining the analytical analysis of multiple pharmacology assays along with cryo-EM structures and GaMD simulations. Overall, these results provide a framework for future mechanistic studies, and ultimately, aid in the discovery, design, and optimization of allosteric drugs as novel therapeutic candidates for clinical progression.

## Acknowledgments

This work was supported by a Wellcome Trust Collaborative Award (201529/Z/16/Z; P.M.S., A.B.T., A.C.), the National Health and Medical Research Council of Australia (1055134, 1150083, and 1138448), the Australian Research Council (DE170100152, DP190102950, and IC200100052), and the National Institutes of Health (GM132572). P.M.S. is a Senior Principal Research Fellow (1154434), D.W. a Senior Research Fellow (1155302), D.M.T. an Early Career Research Fellow (1196951), and K.L. a Future Fellow (160100075). R.D. was supported by Takeda Science Foundation 2019 Medical Research Grant and Japan Science and Technology Agency PRESTO (18069571). This work was partially supported by the Monash University Ramaciotti Centre for cryo-electron microscopy and the Monash University MASSIVE high-performance computing facility and supercomputing resources with the XSEDE allocation award TG-MCB180049 and NERSC project M2874. We thank John Tesmer for discussion and the suggestion of calculating DAQ scores.

## Author Contributions

D.M.T, A.C., P.M.S., and A.B.T. designed the overall research; Z.V. and D.M.T designed, expressed, and purified protein samples; Yi-L.L. and A.G. performed negative-stain EM; R.D. performed sample vitrification and cryo-EM imaging; M.J.B, R.D., J.M., and D.M.T processed the EM data; J.M., Z.V., and D.M.T. generated and analysed atomic models; J.W, A.B, and Y.M designed, performed, analysed GaMD simulations, and contributed to writing. V.P., V.N., K.L., W.A.C.B, E.T.W., E.K., G.T., and M.Y., generated DNA constructs and performed pharmacology experiments. D.M.T, C.V., V.P., and K.L. analysed pharmacology data. C.W.L. provided VU0467154. D.W., P.M.S, A.B.T., Y.M., A.C., and D.M.T provided supervision. D.M.T. and A.C. wrote the manuscript with contributions and input from all authors.

## Data and materials availability

All data generated or analysed during this study are included in the manuscript. Structural data has been deposited in the Protein Data Bank (PDB) and Electron Microscopy Data Bank (EMDB) under the following codes:

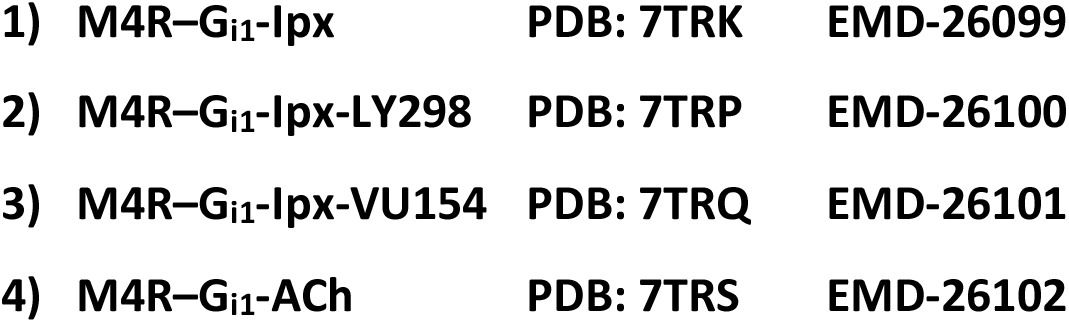

|  |  |  |  |
| --- | --- | --- | --- |
| 737 | **1) M4R-G<sub>i1</sub>-Ipx** | **PDB: 7TRK** | **EMD-26099** |
| 738 | **2) M4R-G<sub>i1</sub>-Ipx-LY298** | **PDB: 7TRP** | **EMD-26100** |
| 739 | **3) M4R-G<sub>i1</sub>-Ipx-VU154** | **PDB: 7TRQ** | **EMD-26101** |
| 740 | **4) M4R-G<sub>i1</sub>-ACh** | **PDB: 7TRS** | **EMD-26102** |

## Materials and Methods

### Key Resources Table

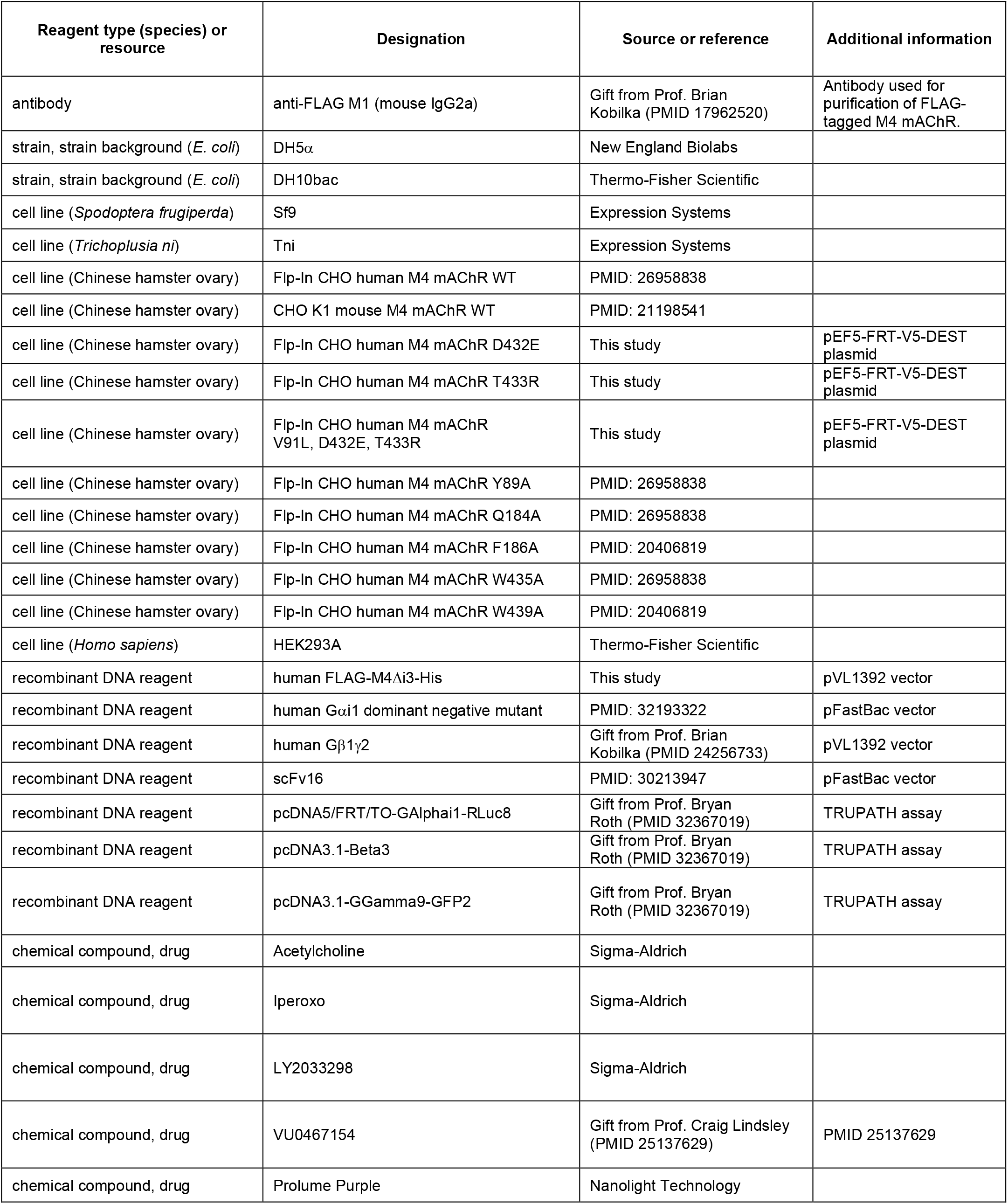
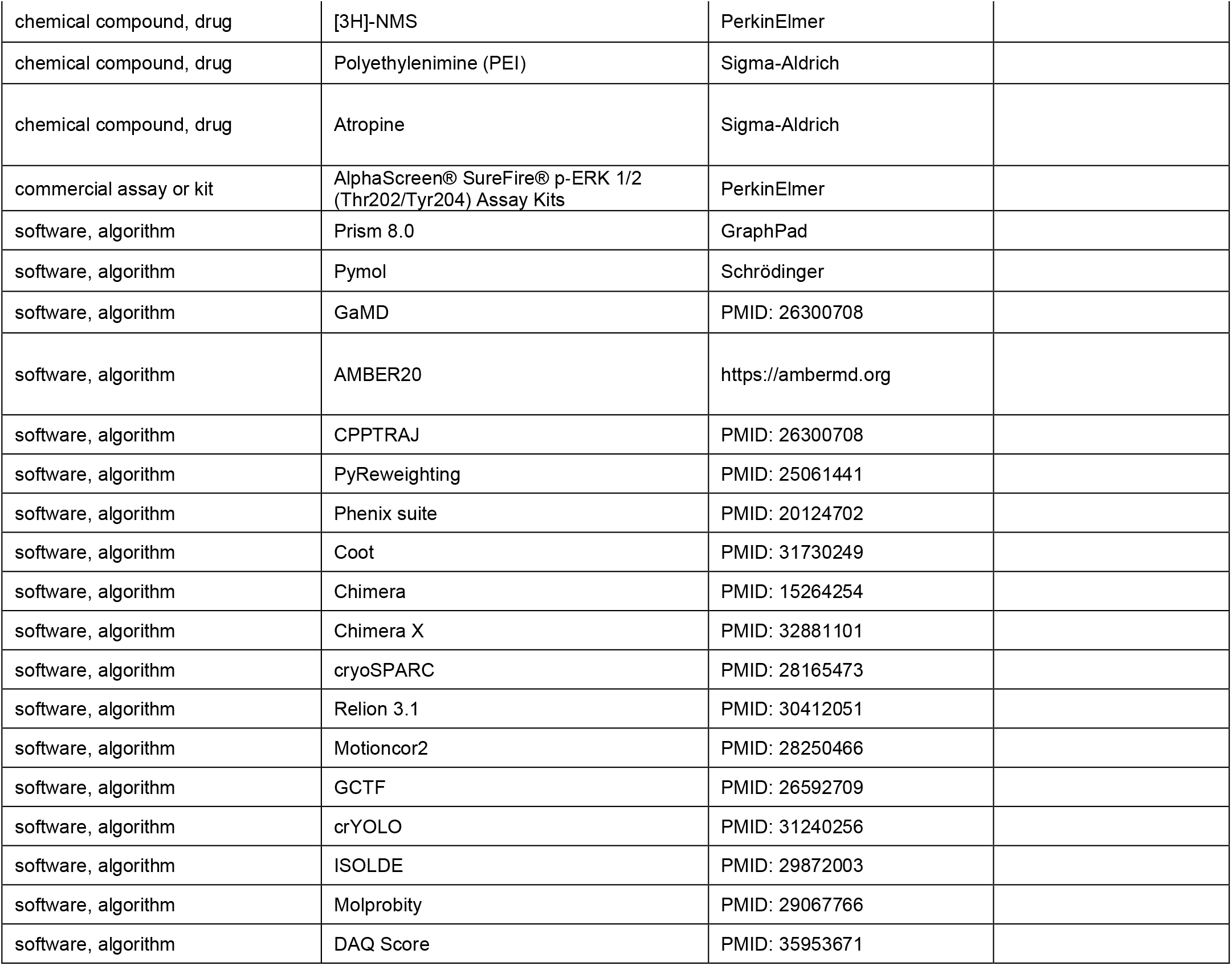

### Bacterial strains

DH5α (New England Biolabs) and DH10bac () *E. coli* cells were grown in LB at 37°C.

### Insect cell culture

Tni and Sf9 cells (Expression Systems) were maintained in ESF-921 media (Expression Systems) at 27°C.

### Mammalian cell culture

Flp-In Chinese hamster ovary (CHO) (Thermo-Fisher Scientific) cells stably expressing human M_4_ mAChR or mutant constructs were maintained in Dulbecco’s modified Eagle’s medium (DMEM, Invitrogen) containing 5% fetal bovine serum (FBS; ThermoTrace) and 0.6 μg/ml of Hygromycin (Roche) in a humidified incubator (37°C, 5% CO_2_, 95% O_2_). HEK293A cells were grown in DMEM supplemented with 5% FBS at 37°C in 5% CO_2_.

### Radioligand binding assays

Flp-In CHO cells stably expressing M_4_ mAChR constructs were seeded at 10,000 cells/well in 96-well white clear bottom isoplates (Greiner Bio-one) and allowed to adhere overnight at 37°C, 5% CO_2_, and 95% O_2_. Saturation binding assay was performed to quantify the receptor expression and equilibrium dissociation constant of the radioligand [^3^H]-NMS (PerkinElmer, specific activity 80 Ci/mmol). Briefly, plates were washed once with phosphate-buffered saline (PBS) and incubated overnight at room temperature (RT) with 0.01-10 nM [^3^H]-NMS in Hanks’s balanced salt solution (HBSS)/10 mM HEPES (pH 7.4) in a final volume of 100 μl. For binding interaction assays, cells were incubated overnight at RT with a specific concentration of [^3^H]-NMS (pK_D_ determined at each receptor in saturation binding) and various concentrations of ACh or iperoxo (Ipx) in the absence or presence of increasing concentrations of each allosteric modulator. In all cases, nonspecific binding was determined by the coaddition of 10 μM atropine (Sigma). The following day, the assays were terminated by washing the plates twice with ice-cold 0.9% NaCl to remove the unbound radioligand. Cells were solubilised in 100 μl per well of Ultima Gold (PerkinElmer), and radioactivity was measured with a MicroBeta plate reader (PerkinElmer).

### G protein activation assay

Upon 60-80% confluence, HEK293A cells were transfected transiently using polyethylenimine (PEI, Polysciences) and 10 ng per well of each of pcDNA3.1-hM4 mAChR (WT or mutant), pcDNA5/FRT/TO-Gα_i1_-RLuc8, pcDNA3.1-β_3_, and pcDNA3.1-Gγ_9_-GFP2 at a ratio of 1:1:1:1 ratio with 40 ng of total DNA per well. Cells were plated at 30,000 cells per well into 96-well Greiner CELLSTAR white-walled plates (Sigma Aldrich). 48 hrs later, cells were washed with 200 μL phosphate buffer saline (PBS) and replaced with 70 μL of 1x HBSS with 10 mM HEPES. Cells were incubated for 30 min at 37°C before addition of 10 μL of 1.3 μM Prolume Purple coelenterazine (Nanolight technology). Cells were further incubated for 10 min at 37C° before BRET measurements were performed on a PHERAstar plate reader (BMG Labtech) using 410/80-nm and 515/30-nm filters. Baseline measurements were taken for 8 min before addition of drugs or vehicle to give a final assay volume of 100 μL and further reading for 30 min. BRET signal was calculated as the ratio of 515/30-nm emission over 410/80-nm emission. The ratio was vehicle corrected using the initial 8 min of baseline measurements and then baseline corrected again using the vehicle-treated wells. Data were normalized using the maximum agonist response to allow for grouping of results using an area under the curve analysis in Prism. Data were analysed at timepoints of 4, 10, and 30 min yielding similar results.

### Phospho-ERK1/2 assay

The level of phosphorylated extracellular signal-regulated protein kinase 1/2 (pERK1/2) was detected using the AlphaScreen^TM^ SureFire Kit (PerkinElmer Life and Analytical Sciences). Briefly, FlpIn CHO cells stably expressing the receptor were seeded into transparent 96-well plates at a density of 20,000 cells/well and grown overnight at 37°C, 5% CO_2_. Cells were washed with PBS and incubated in serum-free DMEM at 37°C for 4 hr to allow FBS-stimulated pERK1/2 levels to subside. Cells were stimulated with increasing concentrations of ACh or Ipx in the absence or presence of increasing concentrations of the allosteric modulator at 37°C for 5 min (the time required to maximally promote ERK phosphorylation for each ligand at each M_4_ mAChR construct in the initial time-course study; data not shown). For all experiments, stimulation with 10% (*v/v*) FBS for 5 min was used as a positive control. The reaction was terminated by the removal of media and lysis of cells with 50 μl of the SureFire lysis buffer (TGR Biosciences). Plates were then agitated for 5 min and 5 μl of the cell lysate was transferred to a white 384-well ProxiPlate (Greiner Bio-one) followed by the addition of 5 μl of the detection buffer (a mixture of activation buffer: reaction buffer: acceptor beads: donor beads at a ratio of 50:200:1:1). Plates were incubated in the dark for 1 hr at 37°C followed by measurement of fluorescence using an Envision plate reader (PerkinElmer) with standard AlphaScreen^TM^ settings. Data were normalised to the maximal response mediated by 10 μM ACh, Ipx or 10% FBS.

### Purification of scFv16

Tni insect cells were infected with scFv16 baculovirus at a density of 4 million cells per mL and harvested at 60 hrs post infection by centrifugation for 10 min at 10,000xg. The supernatant was pH balanced to pH 7.5 by the addition of Tris pH 7.5, and 5 mM CaCl_2_ was added to quench any chelating agents, then left to stir for 1.5 hrs at RT. The supernatant was then centrifuged at 30,000xg for 15 min to remove any precipitates. 5 mL of EDTA resistant Ni resin (Cytivia) was added and incubated for 2 hrs at 4°C while stirring. Resin was collected in a glass column and washed with 20 column volumes (CVs) of high salt buffer (20 mM HEPES pH 7.5, 500 mM NaCl, 20 mM imidazole) followed by 20 CVs of low salt buffer (20 mM HEPES pH 7.5, 100 mM NaCl, 20 mM Imidazole). Protein was then eluted using 8 CV of elution buffer (20 mM HEPES pH 7.5, 100 mM NaCl, 250 mM imidazole) until no more protein was detected using Bradford reagent (Bio-Rad Laboratories). Protein was concentrated using a 10-kDa Amicon filter device (Millipore) and aliquoted into 1 mg aliquots for further use.

### Expression and purification of M_4_R-G_i1_-scFv16 complexes

The human M_4_ mAChR with residues 242 to 387 of the third intracellular loop removed and the N-terminal glycosylation sites (N3, N9, N13) mutated to D was expressed in Sf9 insect cells, and human DNG_αi1_ and His6-tagged human G_β1γ2_ were co-expressed in Tni insect cells. Cell cultures were grown to a density of 4 million cell per ml for Sf9 cells and 3.6 million per ml for Tni cells and then infected with either M_4_ mAChR baculovirus or both G_αi1_ and G_β1γ2_ baculovirus, at a ratio of 1:1. M_4_ mAChR expression was supplemented with 10 mM atropine. Cultures were grown at 27°C and harvested by centrifugation 60-72 hr (48 h for Hi5 cells) post infection. Cells were frozen and stored at −80°C for later use. 1-2 L of the frozen cells were used for each purification.

Cells expressing M_4_ mAChR were thawed at RT and then dounced in the solubilization buffer containing: 20 mM HEPES pH 7.5, 10% glycerol, 750 mM NaCl, 5 mM MgCl_2_, 5 mM CaCl_2_, 0.5% LMNG, 0.02% CHS, 10 µM atropine and cOmplete protease inhibitor cocktail (Roche) until homogenous. The receptor was solubilized for 2 hrs at 4°C while stirring. The insoluble material was removed by centrifugation at 30,000xg for 30 min followed by filtering the supernatant and batch-binding immobilization to M1 anti-flag affinity resin, previously equilibrated with high salt buffer, for 1 hr at RT. The resin with immobilized receptor was then washed using a peristaltic pump for 30 min at 2 ml/min with high salt buffer: 20 mM HEPES pH 7.5, 750 mM NaCl, 5 mM MgCl_2_, 5 mM CaCl_2_, 0.5% lauryl maltose neopentyl glycol (LMNG, Anatrace), 0.02% cholesterol hemisuccinate (CHS, Anatrace) followed by low salt buffer: 20 mM HEPES pH 7.5, 100 mM NaCl, 5 mM MgCl_2_, 5 mM CaCl_2_, 0.5% LMNG, 0.02% CHS and an agonist (5 µM Ipx, 1µM Ipx with 10 µM VU154, or 100 µM ACh). While the receptor was immobilised on anti-FLAG resin, the DNGα_i1_ cell pellet was thawed, dounced, and solubilised in the solubilisation buffer containing: 20 mM HEPES pH 7.5, 100 mM NaCl, 5 mM MgCl_2_, 5 mM CaCl_2_, 0.5% LMNG, 0.02% CHS, apyrase (5 units), and cOmplete Protease Inhibitor Cocktail. DNGα_i1_ was solubilized for 2 hrs at 4°C followed by the centrifugation at 30,000xg for 30 min to remove the insoluble material. Supernatant was filtered through a glass fibre filter (Millipore) and then added to the receptor bound to anti-Flag resin. Apyrase (5 units), scFv16, and agonist (either 1 µM Ipx, 1 µM Ipx with 10 µM VU154, or 100 µM ACh) were added and incubated for 1 hr at RT with gentle mixing. The anti-FLAG resin was then loaded onto a glass column and washed with approximately 20 CVs of washing buffer: 20 mM HEPES pH 7.4, 100 mM NaCl, 5 mM MgCl_2_, 5 mM CaCl_2_, 0.01% LMNG, 0.001% CHS, agonist (1 µM Ipx, 1 µM Ipx with 10 µM VU154, or 100 µM ACh). Complex was eluted with size-exclusion chromatography (SEC) buffer: 20 mM HEPES pH 7.5, 100 mM NaCl, 5 mM MgCl_2_, 0.01% LMNG, 0.001% CHS and agonist (1 µM Ipx, or 1 µM Ipx with 10 µM VU154, or 100 µM ACh) with the addition of 10 mM EGTA and 0.1 mg/mL FLAG peptide. After the elution an additional 1-2 mg of scFv16 was added, and shortly incubated on ice before concentrating using a 100-kDa Amicon filter to a final volume of 500 µL. The sample was filtered using a 0.22 µm filter followed by SEC using a Superdex 200 increase 10/300 column (Cytivia) using SEC buffer. For the ACh- and VU154-Ipx-bound samples, the fractions containing protein were concentrated again and re-run over SEC using a buffer with half the amount of detergent in order to remove empty micelles. Samples were concentrated and flash frozen using liquid nitrogen. In case of the LY298-Ipx-bound sample, the sample was purified with 1 µM Ipx only. After SEC, the sample was then split in half, where one half was incubated with approximately 1.6 µM LY298 at 4°C overnight, and then concentrated and flash frozen in liquid nitrogen.

### EM sample preparation and data acquisition

Samples (3 µL) were applied to glow-discharged Quantifoil R1.2/1.3 Cu/Rh 200 mesh grids (Quantifoil) (M4R-G_i1_-Ipx and M4R-G_i1_-Ipx-LY298) or UltrAuFoil R1.2/1.3 Au 300 mesh grids (Quantifoil) (M4R-G_i1_-Ipx-VU154 and M4R-G_i1_-Ach) and were vitrified on a Vitrobot Mark IV (Thermo Fisher Scientific) set to 4°C and 100 % humidity and 10 s blot time. Data were collected on a Titan Krios G3i 300 kV electron microscope (Themo Fisher Scientific) equipped with GIF Quantum energy filter and K3 detector (Gatan). Data acquisition was performed in EFTEM NanoProbe mode with a 50 µM C2 aperture at an indicated magnification of ×105,000 with zero-loss slit width of 25 eV. The data were collected automatically with homemade scripts for SerialEM performing a 9-hole beam-image shift acquisition scheme with one exposure in the centre of each hole. Experimental parameters specific to each collected data set is listed in Table S2.

### Image processing

Specific details for the processing of each cryo-EM data set are shown in Figure S1. Image frames for each movie were motion corrected using MotionCor2 (Zheng et al., 2017) and contrast transfer function (CTF)-estimated using GCTF (Zhang, 2016). Particles were picked from corrected micrographs using crYOLO (Wagner et al., 2019) or RELION-3.1 software (Zivanov et al., 2018) followed by reference-free 2D and 3D classifications. Particles within bad classes were removed and remaining particles subjected to further analysis. Resulting particles were subjected to Bayesian polishing, CTF refinement, 3D auto-refinement in RELION, followed by another round of 3D classification and 3D refinement that yielded the final maps (Zivanov et al., 2018). Local resolution was determined from RELION using half-reconstructions as input maps. Due to the high degree of conformational flexibility between the receptor and G protein, a further local refinement was performed in cryoSPARC for the ACh-bound M_4_R-complex. A receptor focused map was generated (2.75 Å) which was used to generate a PDB model of the ACh-bound M_4_R.

### Model building and refinement

An initial M_4_R template model was generated from our prior modelling studies of the M_4_ mAChR that was based on an active state M_2_ mAChR structure (PBD: 4MQT) (Kruse et al., 2013). An initial model for dominant negative Gα_i1_Gβ_1_Gγ_2_ was from a structure in complex with Smoothend (PDB: 6OT0) (Qi et al., 2019) and scFv16 from the X-ray crystal structure in complex with heterotrimeric G protein (PDB: 6CRK) (Maeda et al., 2018). Models were fit into EM maps using UCSF Chimera (Pettersen et al., 2004), and then rigid-body-fit using PHENIX (Liebschner et al., 2019), followed by iterative rounds of model rebuilding in Coot (Casañal et al., 2020) and ISOLDE (Croll, 2018), and real-space refinement in PHENIX. Restrains for all ligands were generated from the GRADE server, https://grade.globalphasing.org. Model validation was performed with Molprobity (Williams et al., 2018) and the wwPDB validation server (Berman et al., 2003). Figures were generated using UCSF Chimera (Pettersen et al., 2004), Chimera X (Pettersen et al., 2021), and PyMOL (Schrödinger).

### Cryo-EM 3D variability analysis

3D variability analysis (3DVAR) was performed to access and visualize the dynamics within the cryo-EM datasets of the M_4_ mAChR complexes, as previously described using cryoSPARC (Punjani and Fleet, 2021). The polished particle stacks were imported into cryoSPARC, followed by 2D classification and 3D refinement using the respective low pass filtered RELION consensus maps as an initial model. 3DVA was analysed in three components with 20 volume frames of data per component of motion. Output files were visualized using UCSF Chimera (Pettersen et al., 2004).

#### Gaussian accelerated molecular dynamics (GaMD)

GaMD enhances the conformational sampling of biomolecules by adding a harmonic boost potential to reduce the system energy barriers (Miao et al., 2015). When the system potential 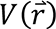 is lower than a reference energy E, the modified potential 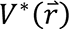 of the system is calculated as:

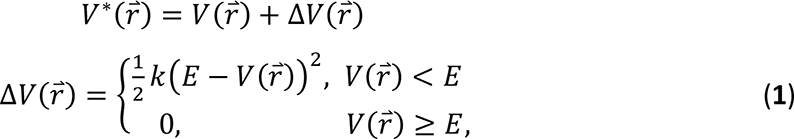

Where k is the harmonic force constant. The two adjustable parameters E and k are automatically determined on three enhanced sampling principles. First, for any two arbitrary potential values 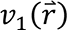 and 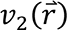 found on the original energy surface, if 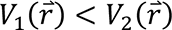, Δ*V* should be a monotonic function that does not change the relative order of the biased potential values; i.e., 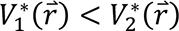. Second, if 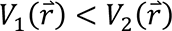, the potential difference observed on the smoothened energy surface should be smaller than that of the original; i.e., 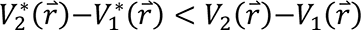. By combining the first two criteria and plugging in the formula of 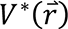 and Δ*V*, we obtain

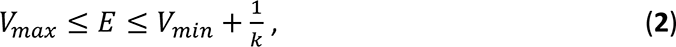

Where *V*_*min*_ and *V*_*max*_ are the system minimum and maximum potential energies. To ensure that Eq. 2 is valid, *k* has to satisfy: *k* ≤ 1/(*V*_*max*_ − *V*_*min*_). Let us define: *k* = *k*_0_ · 1/(*V*_*max*_ − *V*_*min*_), then 0 < *k*_0_ ≤ 1. Third, the standard deviation (SD) of Δ*V* needs to be small enough (i.e. narrow distribution) to ensure accurate reweighting using cumulant expansion to the second order: *σ*_Δ*V*_ = *k*(*E* − *V*_*avg*_)*σ*_*V*_ ≤ *σ*_0_, where *V*_*avg*_ and *σ*_*V*_ are the average and SD of Δ*V*with *σ*_0_ as a user-specified upper limit (e.g., 10*k*_*B*_*T*) for accurate reweighting. When E is set to the lower bound *E* = *V*_*max*_ according to Eq. 2, *k*_0_ can be calculated as

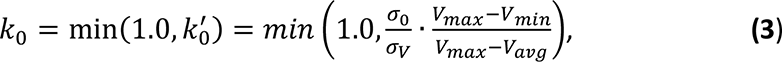

Alternatively, when the threshold energy E is set to its upper bound *E* = *V*_*min*_ + 1/*k*, *k*_0_is set to:

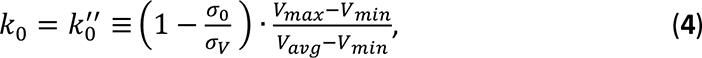

If 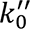 is calculated between 0 and 1. Otherwise, *k*_0_ is calculated using Eq. 3.

#### Energetic Reweighting of GaMD Simulations

For energetic reweighting of GaMD simulations to calculate potential of mean force (PMF), the probability distribution along a reaction coordinate is written as *p*^∗^(*A*) . Given the boost potential Δ*V*(*r*) of each frame, *p*^∗^(*A*) can be reweighted to recover the canonical ensemble distribution *p*(*A*), as:

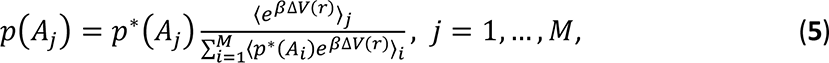

where *M* is the number of bins, *β* = *k*_*B*_*T* and 〈*e*^*β*Δ*V*(*r*)^〉_*j*_ is the ensemble-averaged Boltzmann factor of Δ*V*(*r*) for simulation frames found in the *j*^th^ bin. The ensemble-averaged reweighting factor can be approximated using cumulant expansion:

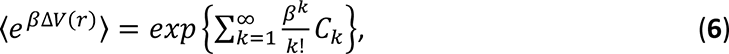

where the first two cumulants are given by:

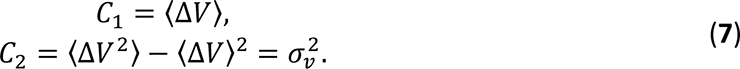

The boost potential obtained from GaMD simulations usually follows near-Gaussian distribution (Miao and McCammon, 2017). Cumulant expansion to the second order thus provides a good approximation for computing the reweighting factor (Miao et al., 2014, 2015). The reweighted free energy *F*(*A*) = −*k*_*B*_*T* ln *p*(*A*) is calculated as:

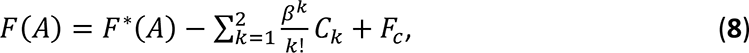

where *F*^∗^(*A*) = −*k*_*B*_*T* ln *p*^∗^(*A*) is the modified free energy obtained from GaMD simulation and *F*_*c*_ is a constant.

#### System Setup

The M_4_R-ACh-G_i1_, M_4_R-Ipx-G_i1_, M_4_R-Ipx-G_i1_-VU154 and M_4_R-Ipx-G_i1_-LY298 cryo-EM structures were used for setting up simulation systems. The scFv16 in the cryo-EM structures was omitted in all simulations. The initial structures of single mutant D432E and T433R mutant of M_4_R-Ipx-G_i1_-VU154 were obtained by mutating the corresponding residues in the M_4_R-Ipx-G_i1_-VU154 cryo-EM structure. The initial structures of M_4_R-ACh-G_i1_-VU154 and M_4_R-ACh-G_i1_-LY298 were obtained from M_4_R-Ipx-G_i1_-VU154 and M_4_R-Ipx-G_i1_-LY298 cryo-EM structures by replacing Ipx with ACh through alignment of receptors to the M4R-ACh-G_i1_ cryo-EM structure. The initial structures of M_4_R-G_i1_-VU154 and M_4_R-G_i1_-LY298 were obtained by removing the corresponding Ipx agonist from the M_4_R-Ipx-G_i1_-VU154 and M_4_R-Ipx-G_i1_-LY298 cryo-EM structures. The initial structures of M_4_R-VU154 and M_4_R-LY298 were obtained by removing the corresponding Ipxagonist and G_i1_ protein from the M_4_R-Ipx-G_i1_-VU154 and M_4_R-Ipx-G_i1_-LY298 cryo-EM structures. According to previous findings, intracellular loop (ICL) 3 is highly flexible and removal of ICL3 does not appear to affect GPCR function (Dror et al., 2011, 2015). The ICL3 was thus omitted as in the current GaMD simulations. Similar as previous study, helical domains of the G_i1_ protein missing in the cryo-EM structures were not included in the simulation models. This was based on earlier simulation of the β_2_AR-G_s_ complex, which showed that the helical domain fluctuated substantially (Dror et al., 2015). All chain termini were capped with neutral groups (acetyl and methylamide). All the disulphide bonds in the complexes (i.e., Cys108^3.25^-Cys185^45×50^ and Cys426^ECL3^-Cys429^ECL3^ in the M4R) that were resolved in the cryo-EM structures were maintained in the simulations. Using the *psfgen* plugin in VMD (Humphrey et al., 1996), missing atoms in protein residues were added and all protein residues were set to the standard CHARMM protonation states at neutral pH. For each of the complex systems, the receptor was inserted into a palmitoyl-oleoyl-phosphatidyl-choline (POPC) bilayer with all overlapping lipid molecules removed using the membrane plugin in VMD. The system charges were then neutralized at 0.15M NaCl using the *solvate* plugin in VMD (Humphrey et al., 1996). The simulation systems were summarized in Table S3.

#### Simulation Protocol

The CHARMM36M parameter set (Huang et al., 2017; Klauda et al., 2010; Vanommeslaeghe and MacKerell, 2015) was used for the M_4_ mAChRs, G_i1_ proteins, and POPC lipids. Force field parameters of agonists ACh and Ipx, PAMs LY298 and VU154 were obtained from the CHARMM ParamChem web server (Vanommeslaeghe and MacKerell, 2012; Vanommeslaeghe et al., 2012). Force field parameters with high penalty were optimized with FFParm (Kumar et al., 2020). GaMD simulations of these systems followed a similar protocol used in previous studies of GPCRs (Draper-Joyce et al., 2021; Miao and McCammon, 2016, 2018). For each of the complex systems, initial energy minimization, thermalization, and 20ns cMD equilibration were performed using NAMD2.12 (Phillips et al., 2005). A cutoff distance of 12 Å was used for the van der Waals and short-range electrostatic interactions and the long-range electrostatic interactions were computed with the particle-mesh Ewald summation method (Darden et al., 1993). A 2-fs integration time step was used for all MD simulations and a multiple-time-stepping algorithm was used with bonded and short-range non-bonded interactions computed every time step and long-range electrostatic interactions every two-time steps. The SHAKE algorithm (Ryckaert et al., 1977) was applied to all hydrogen-containing bonds. The NAMD simulation started with equilibration of the lipid tails. With all other atoms fixed, the lipid tails were energy minimized for 1,000 steps using the conjugate gradient algorithm and melted with a constant number, volume, and temperature (NVT) run for 0.5 ns at 310 K. The twelve systems were further equilibrated using a constant number, pressure, and temperature (NPT) run at 1 atm and 310 K for 10 ns with 5 kcal/(mol· Å^2^) harmonic position restraints applied to the protein and ligand atoms. Final equilibration of each system was performed using a NPT run at 1 atm pressure and 310 K for 0.5 ns with all atoms unrestrained. After energy minimization and system equilibration, conventional MD simulations were performed on each system for 20 ns at 1 atm pressure and 310 K with a constant ratio constraint applied on the lipid bilayer in the X-Y plane.

With the NAMD output structure, along with the system topology and CHARMM36M force field files, the *ParmEd* tool in the AMBER package was used to convert the simulation files into the AMBER format. The GaMD module implemented in the GPU version of AMBER20 (Case et al. 2020) was then applied to perform the GaMD simulation. GaMD simulations of systems with G_i1_ protein (M_4_R-ACh-G_i1_, M_4_R-Ipx-G_i1_, M_4_R-Ipx-G_i1_-VU154, M_4_R-Ipx-G_i1_-LY298, M_4_R-ACh-G_i1_-VU154, M_4_R-ACh-G_i1_-LY298, single mutant D432E and T433R mutants of M_4_R-Ipx-G_i1_-VU154) included a 8-ns short cMD simulation used to collect the potential statistics for calculating GaMD acceleration parameters, a 48-ns equilibration after adding the boost potential, and finally three independent 500-ns GaMD production simulations with randomized initial atomic velocities. The average and SD of the system potential energies were calculated every 800,000 steps (1.6 ns). GaMD simulations of M_4_R-VU154 and M_4_R-LY298 included a 2.4-ns short cMD simulation used to collect the potential statistics for calculating GaMD acceleration parameters, a 48-ns equilibration after adding the boost potential, and finally three independent 1000-ns GaMD production simulations with randomized initial atomic velocities. The average and SD of the system potential energies were calculated every 240,000 steps (0.48 ns). All GaMD simulations were run at the “dual-boost” level by setting the reference energy to the lower bound. One boost potential is applied to the dihedral energetic term and the other to the total potential energetic term. The upper limit of the boost potential SD, σ_0_ was set to 6.0 kcal/mol for both the dihedral and the total potential energetic terms. Similar temperature and pressure parameters were used as in the NAMD simulations.

#### Simulation Analysis

CPPTRAJ (Roe and Cheatham, 2013) and VMD (Humphrey et al., 1996) were used to analyze the GaMD simulations. The root-mean square deviations (RMSDs) of the agonist ACh and Ipx, PAM VU154 and LY298 relative to the simulation starting structures, the interactions between receptor and agonists/PAMs, distances between the receptor TM3 and TM6 intracellular ends were selected as reaction coordinates. Particularly, distances were calculated between the Cα atoms of residues Arg^3.50^ and Thr^6.30^, N atom of residue N117^3.37^ and carbon atom (C5) in the acetyl group of ACh or oxygen atom (O09) in the ether bond of Ipx, NE1 atom of residue W164^4.67^ and carbon atom (C5) in the acetyl group of ACh or oxygen atom (O09) in the ether bond of Ipx, indole ring of residue W413^6.48^ and acetyl group of ACh or heterocyclic isoazoline group of Ipx, OH atom of residue Y89^2.61^ and oxygen atom in the amide group of VU154/LY298, benzene ring of residue F186^45.51^ and aromatic core of the PAMs VU154/LY298, OH atom of residue Y439^7.39^ and nitrogen atoms in the amine group of the PAMs VU154/LY298, CD atom of residue Q184^45.49^ and nitrogen atom in the amide group of VU154/LY298, CG atom of residue N423^6.58^ and chlorine atom in PAM LY298, OH atom of residue Y92^2.64^ and nitrogen atom in the amide group of VU154, OG1 atom of residue T433^7.33^ and sulfur atom in the trifluoromethylsulfonyl group of VU154. In addition, the χ2 angle of residue W413^6.48^ and W435^7.35^ were calculated. Time courses of these reaction coordinates obtained from the GaMD simulation were plotted in Figures 4,5,6,7, S4**, and** S7. The PyReweighting (Miao et al., 2014) toolkit was applied to reweight GaMD simulations to recover the original free energy or potential of mean force (PMF) profiles of the simulation systems. PMF profiles were computed using the combined trajectories from all the three independent 500 ns GaMD simulations for each system. A bin size of 1.0 Å was used for RMSD. The cutoff was set to 500 frames for 2D PMF calculations. The 2D PMF profiles were obtained for wildtype M_4_R-Ipx-G_i1_-LY298, M_4_R-Ipx-G_i1_-VU154, and the D432E and T433R single mutants of the M_4_R-Ipx-G_i1_-VU154 system regarding the RMSDs of the agonist Ipx and the RMSDs of the PAMs relative to the cryo-EM conformation (Fig S7).

### Data analysis

All pharmacological data was fit using GraphPad Prism 9.2.0. Saturation binding experiments to determine B_max_ and pK_d_ values were determined as previously described (Leach et al., 2011; Nawaratne et al., 2010; Thal et al., 2016). Detailed equations and analysis details can be found in **Appendix 1**. Interaction inhibition binding curves between [^3^H]-NMS, agonists (ACh or Ipx), and PAMs (LY298 or VU154) were analysed using the allosteric ternary complex model to calculate binding affinity values for each ligand (pK_A_ – for ACh/Ipx and pK_B_ for LY298/VU154) and the degree of binding modulation between agonist and PAM (log α) (Christopoulos and Kenakin, 2002). The pK_B_ values for LY298 and VU154 were determined from global fits of the ACh and Ipx curves to generate one pK_B_ value per ligand (Ehlert, 1988; Leach et al., 2011; Nawaratne et al., 2010; Thal et al., 2016). All pERK1/2 and TruPath assays were analyzed using the operational model allosterism and agonism to determine values of orthosteric (τ_A_) or allosteric efficacy (τ_B_) and the functional modulation (log αβ) between the agonists and PAMs (Leach et al., 2011; Nawaratne et al., 2010). Binding affinities of the agonists and the PAMs were fixed to values determined from equilibrium binding assays. The τ_B_ values for LY298 and VU154 were determined from global fits of the ACh and Ipx curves (when possible) to generate one value per ligand. For comparison between WT human M_4_ mAChR and other M_4_ mAChR constructs the log τ values were corrected (denoted log τ_C_) by normalizing to B_max_ values from saturation binding experiments (Leach et al., 2011; Nawaratne et al., 2010; Thal et al., 2016). All affinity, potency, and cooperativity values were estimated as logarithms, and statistical analysis between WT and mutant M_4_ mAChR were determined by one-way ANOVA using a Dunnett’s post-hoc test with a value of P < 0.05 considered as significant in this study.

## Supplemental Figures

**Figure S1.**
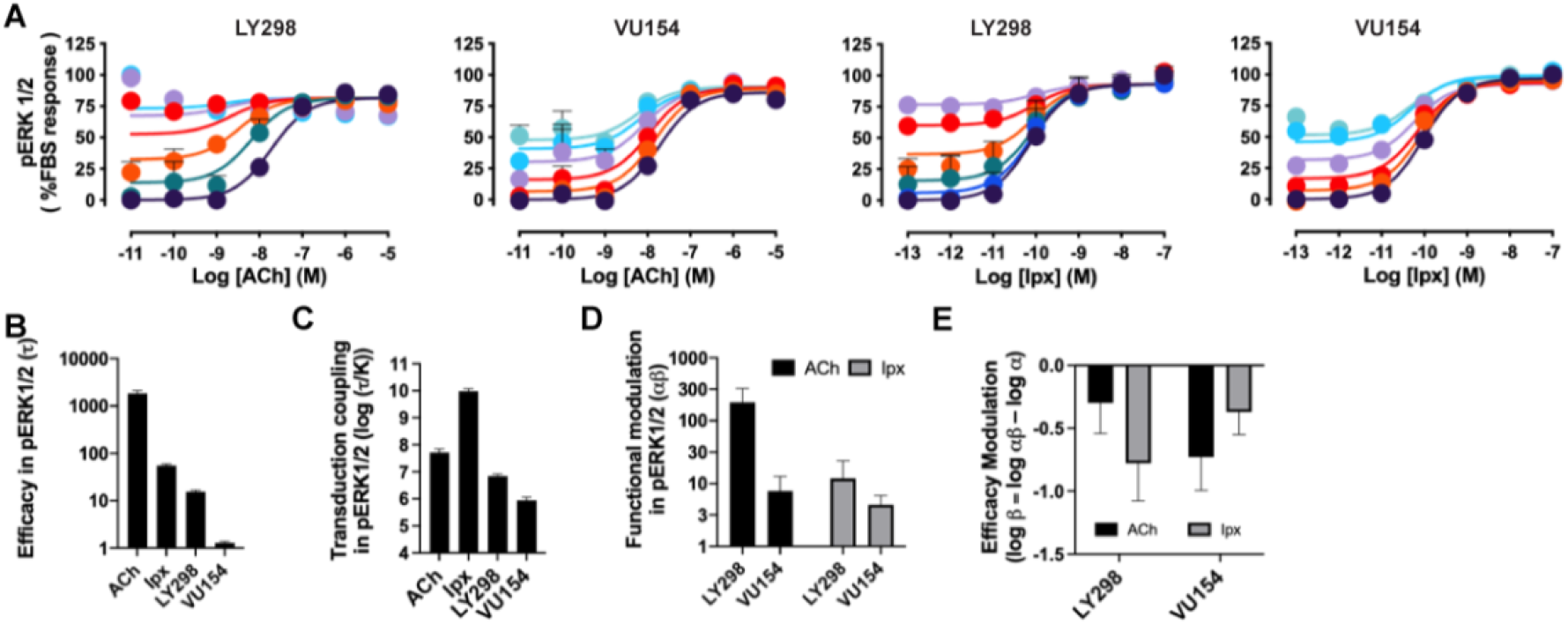
Pharmacological characterization of the PAMs, LY298 and VU154, with ACh and Ipx in pERK1/2 signaling assays, Related to Figure 2. (**A**) Concentration response curves of interactions between the orthosteric and allosteric ligands at the human M_4_ mAChR in the pERK1/2 signaling assay.(**B-E**) Quantification of data from (**A**) and (Figure 2A) to calculate (**B**) the signaling efficacy (τ_A_ and τ_B_) and (**C**) the transduction coupling coefficients (log (τ/K)) of each ligand, (**D**) the functional cooperativity (αβ) between ligands, and (**E**) the efficacy modulation (β) between ligands. All data are mean ± SEM of 3 or more independent experiments performed in duplicate or triplicate with the pharmacological parameters determined from a global fit of the data. The error in (**E**) was propagated using the square root of the sum of the squares. See Table S1.

**Figure S2.**
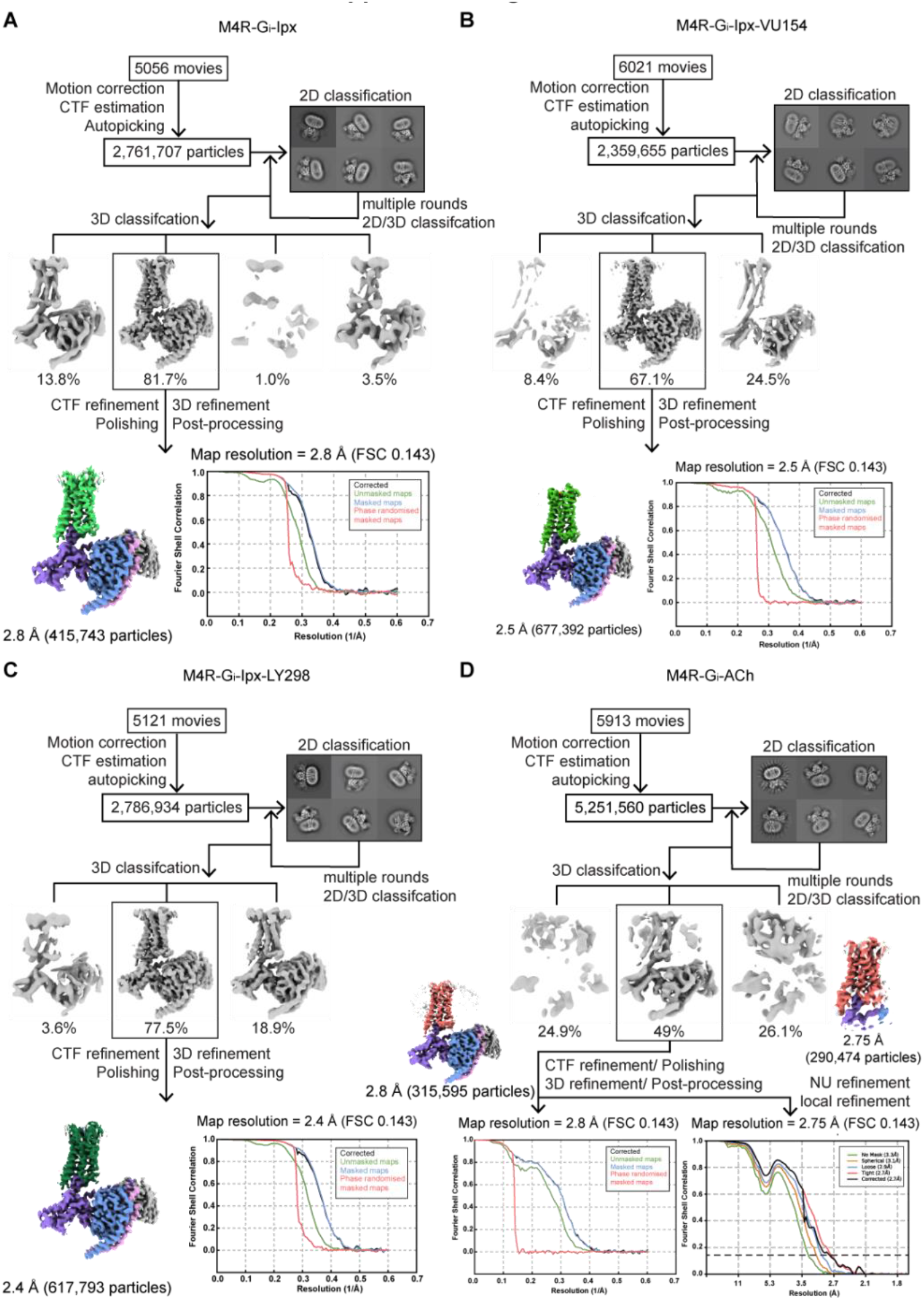
Cryo-EM data processing and analysis, Related to Figure 3. (**A-D**) Flow chart of cryo-EM data processing of the (**A**) Ipx-, (**B**) VU154-Ipx-, (**C**) LY298-Ipx-, and (**D**) ACh-bound M_4_ mAChR complexes with G_i1_-scFv16 including particle selections, 2D and 3D classifications, EM density map, and the Fourier shell correlation (FSC) curves.

**Figure S3.**
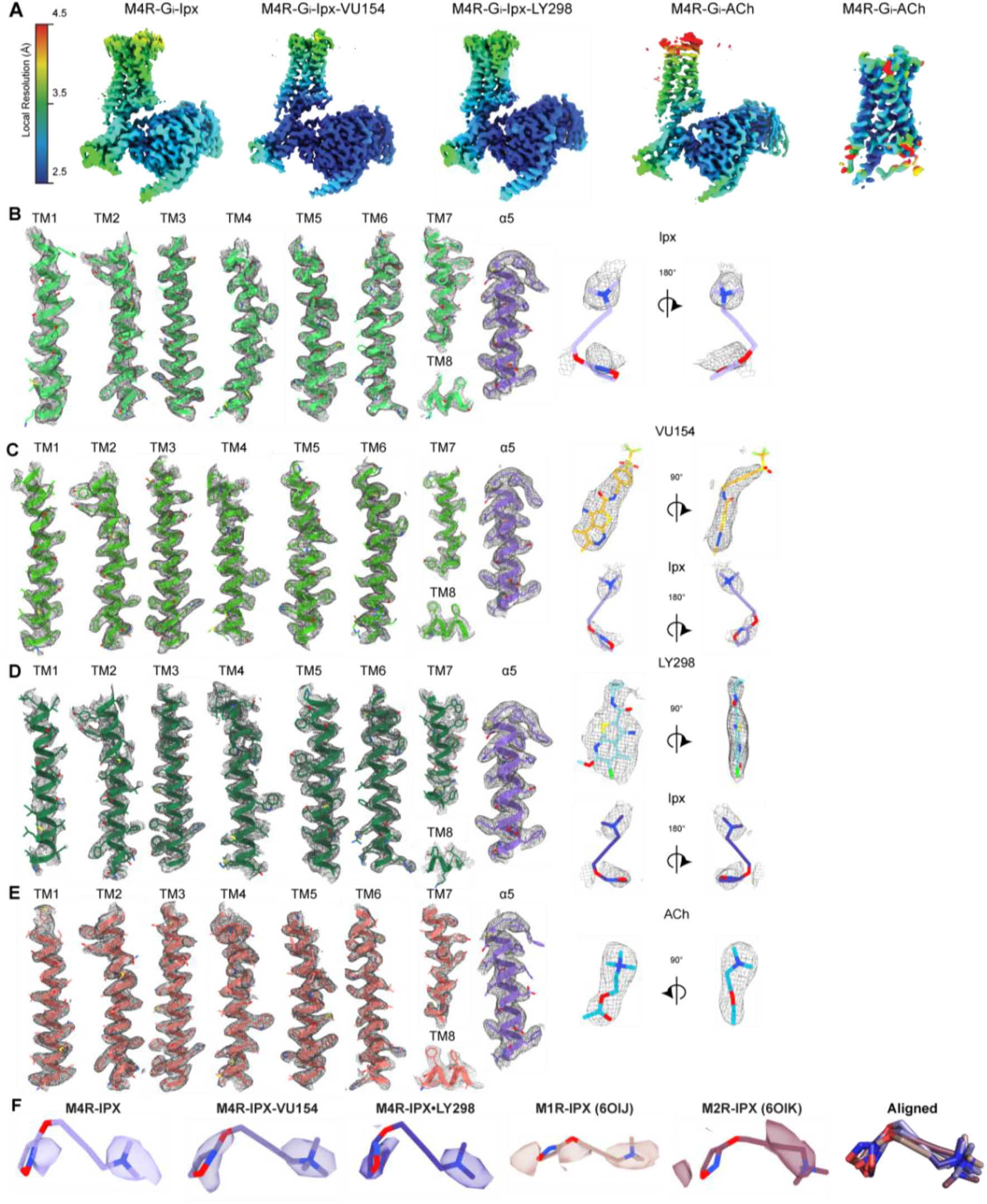
Cryo-EM density maps, Related to Figures 3–4. (**A**) EM maps coloured by local resolution. (**B-E**) Representative EM density and modelling for the 7 transmembrane (TM) helices, the C-terminus of Gα_i1_, and ligands for the (**B**) Ipx-, (**C**) VU154-Ipx-, (**D**) LY298-Ipx-, and (**E**) ACh-bound M_4_ mAChR complexes. (**F**) Comparison of the EM-density of Ipx from other mAChR structures with included Protein Data Bank (PDB) accession codes.

**Figure S4.**
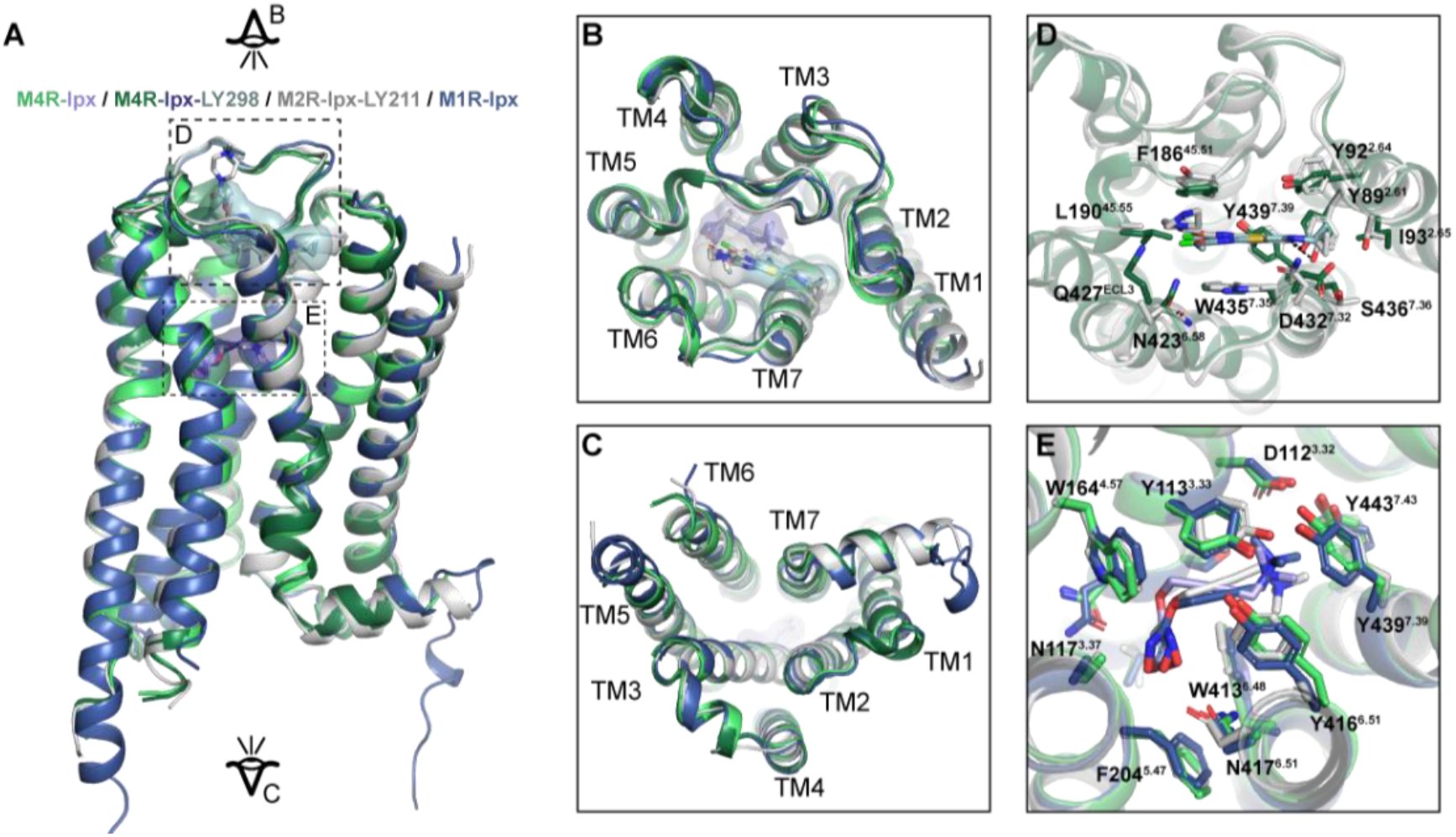
Comparison of active state mAChR structures. (**A**) Comparison of the Ipx- and LY298-Ipx-bound M_4_ mAChR structures to the prior structures of Ipx-bound M_1_ mAChR and LY2119620-Ipx-bound M_2_ mAChR cryo-EM structures. Protein Data Bank (PDB) accession codes for the M_1_ mAChR (PDB: 6OIJ) and the M_2_ mAChR (PDB: 6OIK). (**B,C**) Views from the (**B**) extracellular and (**C**) intracellular surfaces. (**D**) Comparison of the binding pose of LY2119620 at the M_2_ mAChR and LY2033298 at the M_4_ mAChR. (**E**) Comparison of the Ipx binding site residues.

**Figure S5.**
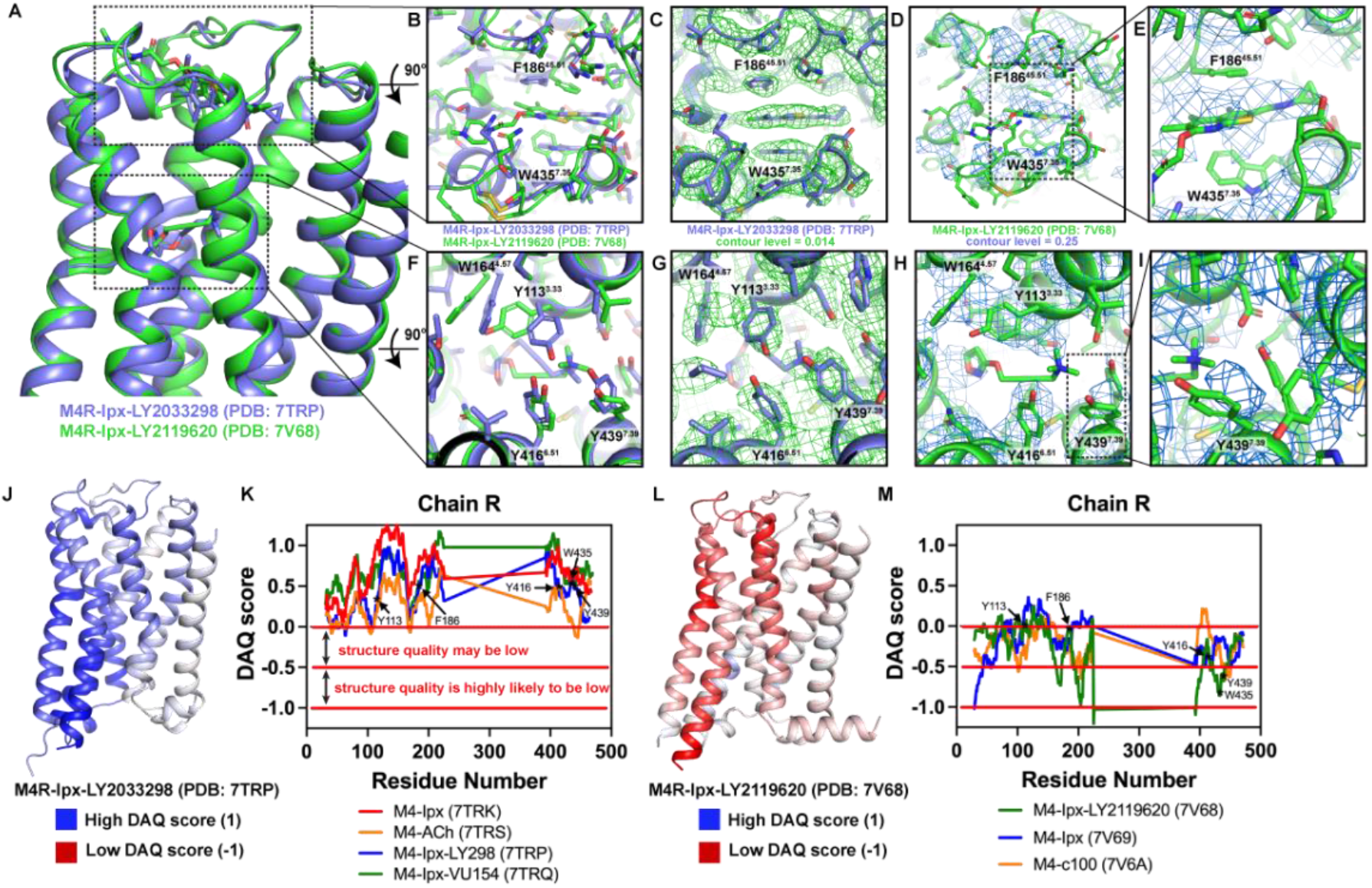
Comparison of active state M_4_ mAChR structures. (**A**) Comparison of LY298-Ipx bound M_4_ mAChR structure (PDB: 7TRP, coloured blue) to the LY2119620-Ipx bound M_4_ mAChR structure (PDB: 7V68, coloured green) (Wang et al., 2022). (**B-E**) View of the allosteric binding site from the top of the receptor. (**B**) Comparison of key allosteric residues F186^45.51^ and W435^7.35^ showing different positions of the residues between M_4_ mAChR structures. (**C**) Overlay of the EM map (EMD-26100, coloured green) onto the LY298-Ipx bound M_4_ mAChR structure contoured at 0.014 in Pymol. (**D-E**) Overlay of the EM map (EMD-31738, coloured blue) onto the LY2119620-Ipx bound M_4_ mAChR structure contoured at 0.25 in Pymol with a close-up of the allosteric residues in (**E**) showing a lack of clear density and mismodelled residues. (**F-I**) View of the orthosteric binding site from the top of the receptor. (**F**) Comparison of key orthosteric binding site residues. (**G**) Related to (**C**) with view from orthosteric site. (**H-I**) Related to (**D**) with view from orthosteric site with a close-up (**I**) of orthosteric residues that are mismodelled or lacking clear density. (**J-M**) DAQ-score provides an estimation of the local quality of protein models from cryo-EM maps on a per residue basis. DAQ-scores were determined from the DAQ web server using the recommended default settings (Terashi et al., 2022).(**J,L**) DAQ scores from the analysis of (J) the LY298-Ipx-M_4_R-G_i1_ complex and (L) the LY2119620-Ipx-M_4_R-G_i1_ complex mapped onto the cartoon of the receptor chain and colour coded by score. A DAQ-score that is positive (coloured blue at values of 1) indicates a correct assignment. A DAQ-score near 0 (colored white) indicates a position in the map that lacks a distinct density pattern for the assigned amino acid. DAQ-scores less than 0 (colored red at −1) indicate a position that could be misassigned or poorly fit. (**K**) DAQ scores for all four M_4_ mAChR structures reported in this manuscript with DAQ scores of each Cα atom plotted for each residue. Key orthosteric and allosteric residues are denoted by asterisks. Nearly every residue has a value above 0. (**M**) Similar to K, but for all three M_4_ mAChR structures reported in (Wang et al., 2022). Very few residues have a score above 0, indicating potential issues with the model and maps.

**Figure S6.**
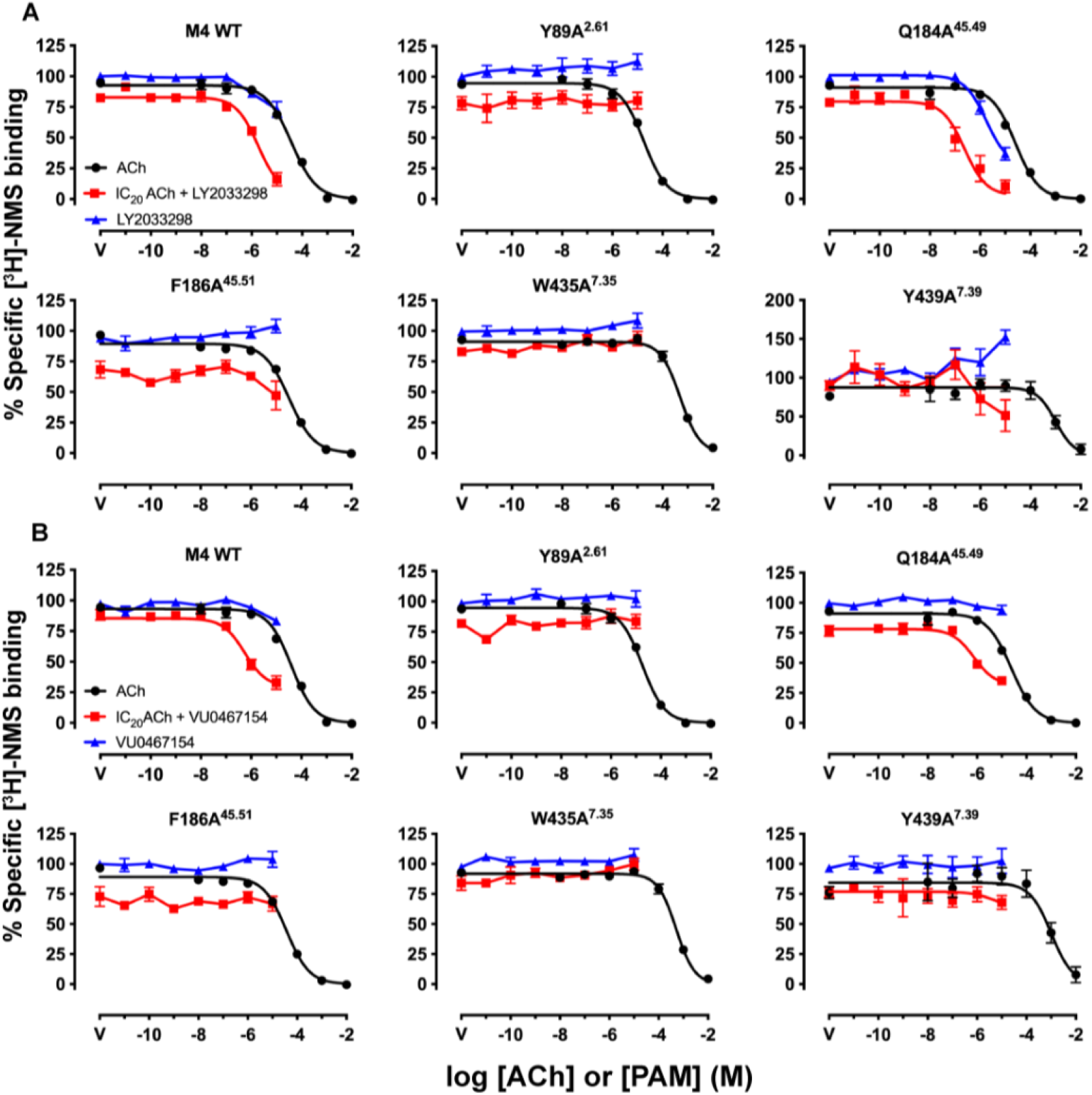
Key residues for the binding of LY298 and VU154 at the human M4 mAChR, Related to Figure 4. (**A,B**) Competition binding with a fixed concentration of [^3^H]-NMS and increasing concentrations of ACh (black circles), (**A**) LY298 or (**B**) VU154 (blue circles), and LY298 or VU154 in the presence of an IC_20_ concentration of ACh (red squares). Curves drawn through the points represent a global fit of an extended ternary complex model. Data points represent the mean ± SEM of 3 or more independent experiments performed in duplicate. Similar data were observed for competition binding with Ipx instead of ACh. See Table S4.

**Figure S7.**
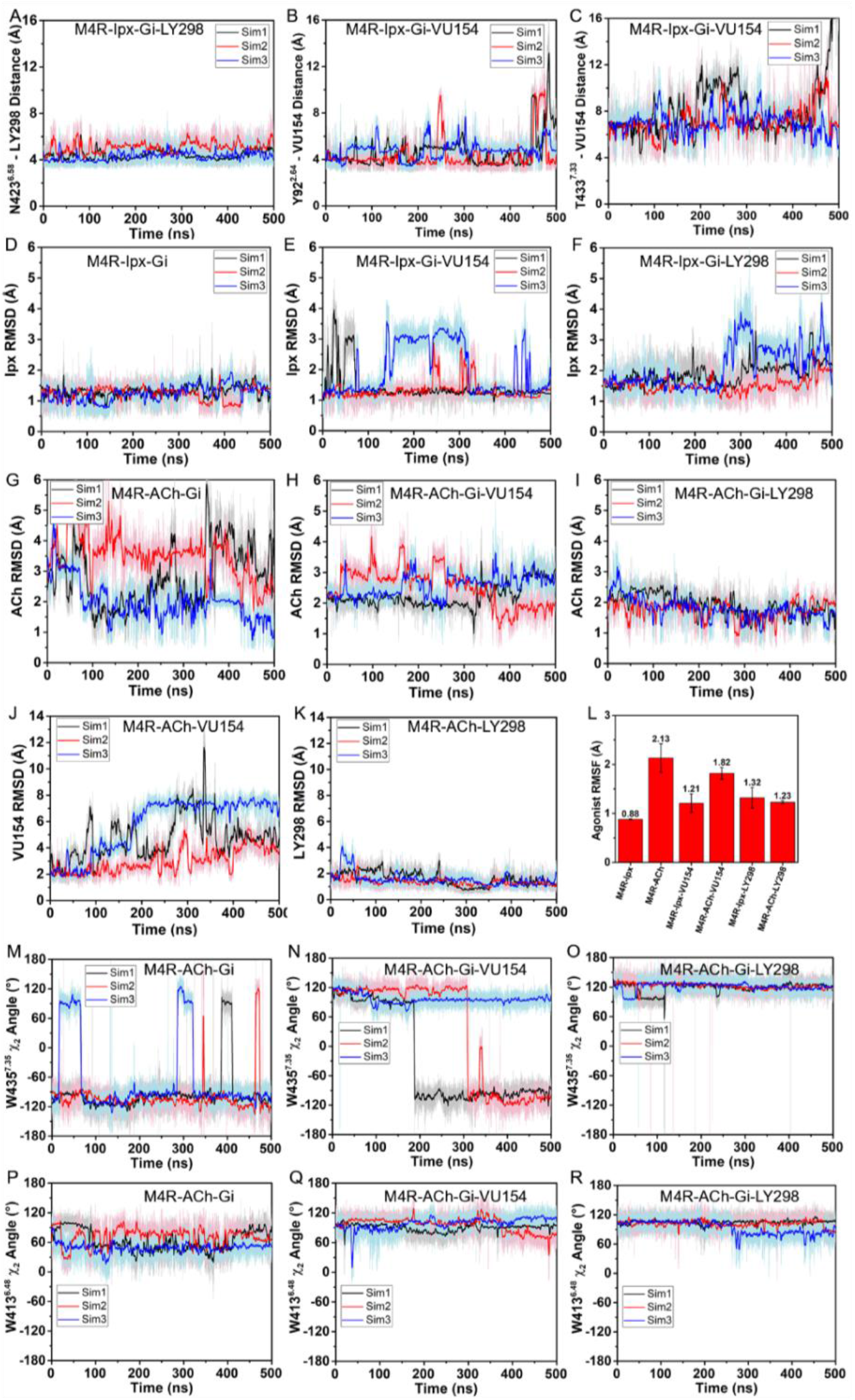
GaMD Simulations, Related to Figures 4–6. (**A-R**) Time courses from GaMD simulations, each performed with 3 separate replicates. Individual replicate simulations are illustrated with different colours. The heading of each plot refers to the specific model used in the simulations. See Table S2. (**A**) Distance between N423^6.58^ and the fluorine atom of LY298 from GAMD simulations of the LY298-Ipx-M4R-G_i1_ structure. (**B,C**) Distance between (**B**) Y92^2.64^ and (**C**) T433^7.33^ to VU154 from GAMD simulations of the VU154-Ipx-M4R-G_i1_ structure. (**D-F**) RMSDs of Ipx from simulations of the cryo-EM structures. (**G-I**) RMSDs of ACh from simulations of the (**G**) cryo-EM structure or (**H,I**) PAM docked models. (**J,K**) RMSDs of VU154 and LY298 from the ACh-bound M4 mAChR simulations. (**L**) Bar graph of the root mean fluctuations of the agonists Ipx or ACh across the GaMD simulations of the M_4_-G_i1_ complexes with or without the PAMs. Values shown are mean ± SEM, n=3. (**M-R**) Time course of the ACh-bound M_4_-G_i1_ simulations illustrating variances in the (**M-O**) W435^7.35^ χ_2_ angle and (**P-R**) the W413^6.48^ χ_2_ angle.

**Figure S8.**
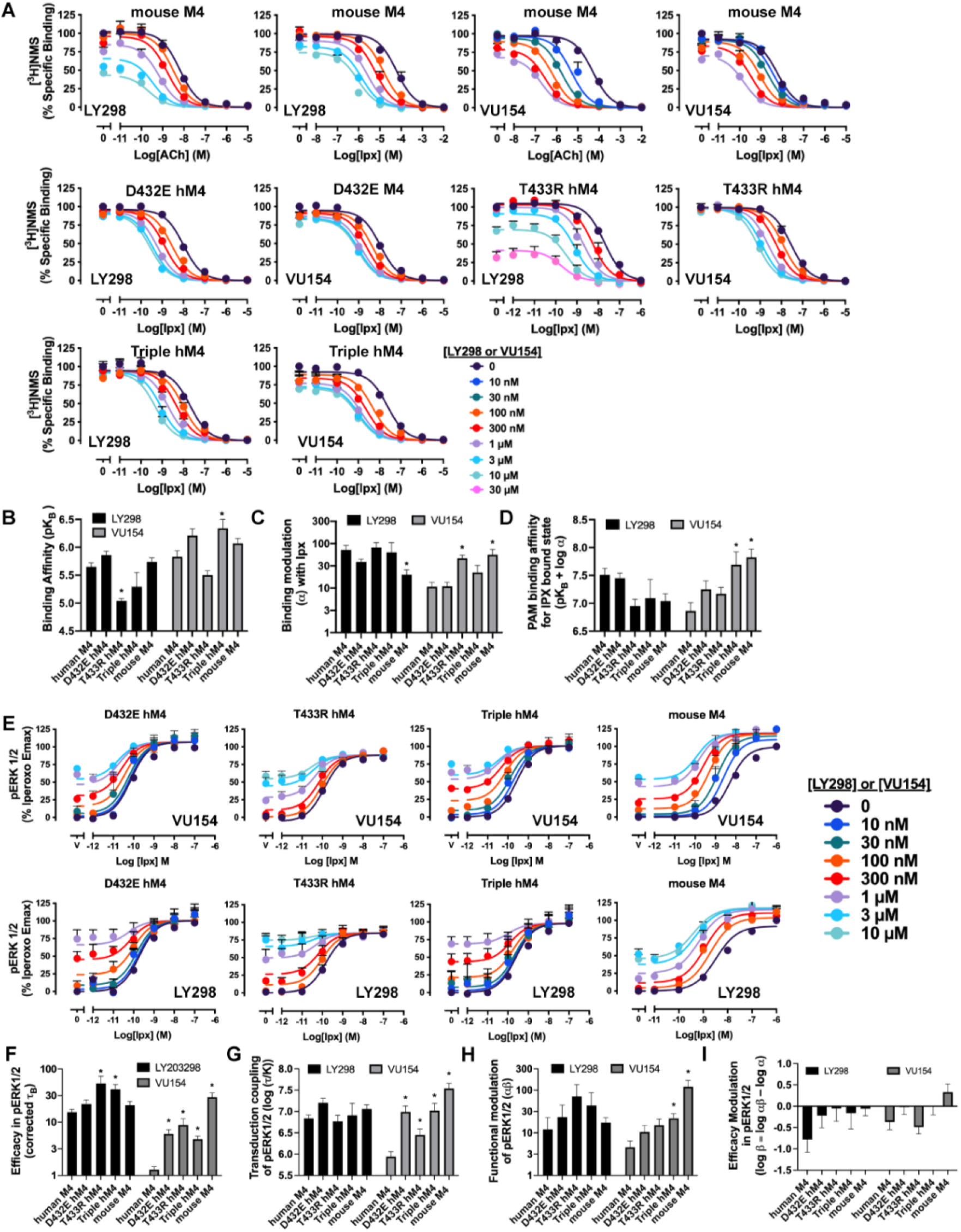
Concentration response curves, Related to Figure 7. (**A**) Concentration response curves the orthosteric and allosteric ligands in [^3^H]-NMS binding assays at the mouse M_4_ mAChR, D432E, T433R, and the V91L, D432E, T433R triple mutant of the human M_4_ mAChR. (**B-D**) Quantification of data from (**A**) to calculate (**B**) equilibrium binding affinities (pK_B_) of the PAMs, (**C**) the degree of binding modulation (α) between Ipx and PAMs, and the modified affinities (**D**) αK_B_. See Table S4. (**E**) Concentration response curves of an interaction between ACh and LY298 in pERK1/2 at the mouse M_4_ mAChR, D432E, T433R, and the V91L, D432E, T433R triple mutant of the human M_4_ mAChR. (**F-I**) Quantification of data from (**A,E**) to calculate (**F**) the signaling efficacy (τ_A_ and τ_B_) and (**G**) the transduction coupling coefficients (log (τ/K)) of each ligand, (**H**) the functional cooperativity (αβ) between ligands, and (**I**) the efficacy modulation (β) between ligands. All data are mean ± SEM of 3 or more independent experiments performed in duplicate or triplicate with the pharmacological parameters determined from a global fit of the data. The error in (**D,I**) was propagated using the square root of the sum of the squares. *Indicates statistical significance (p < 0.05) relative to WT as determined by a one-way ANOVA with a Dunnett’s post-hoc test. See Table S1.

**Figure S9.**
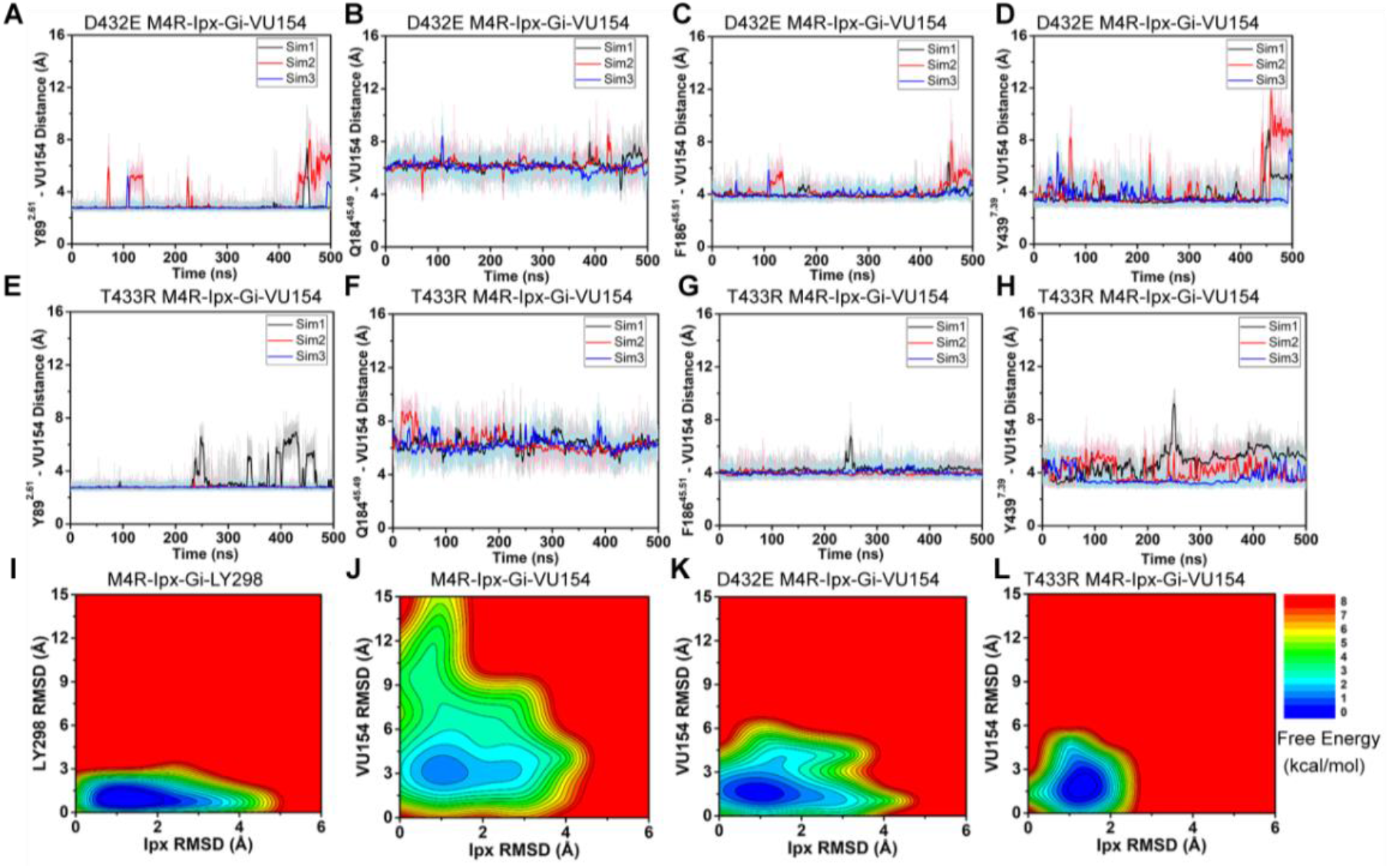
GaMD Simulations of D432E and T433R human M4 mAChR mutants, Related to Figure 7. (**A-H**) Time courses obtained from GaMD simulations of the (**E-H**) D432E and (**I-L**) T433R mutant M_4_R-Ipx-G_i1_-VU154 systems with (**A,E**) Y89^2.61^ – VU154 distance, (**B,F**) Q184^45.49^ – VU154 distance, (**C,G**) F186^45.51^ – VU154 distance, and (**D,H**) Y439^7.39^ – VU154 distance. (**I-L**) 2D free energy profile of the RMSDs of LY298 and VU154 with Ipx. See Table S3.

**Table S1.**
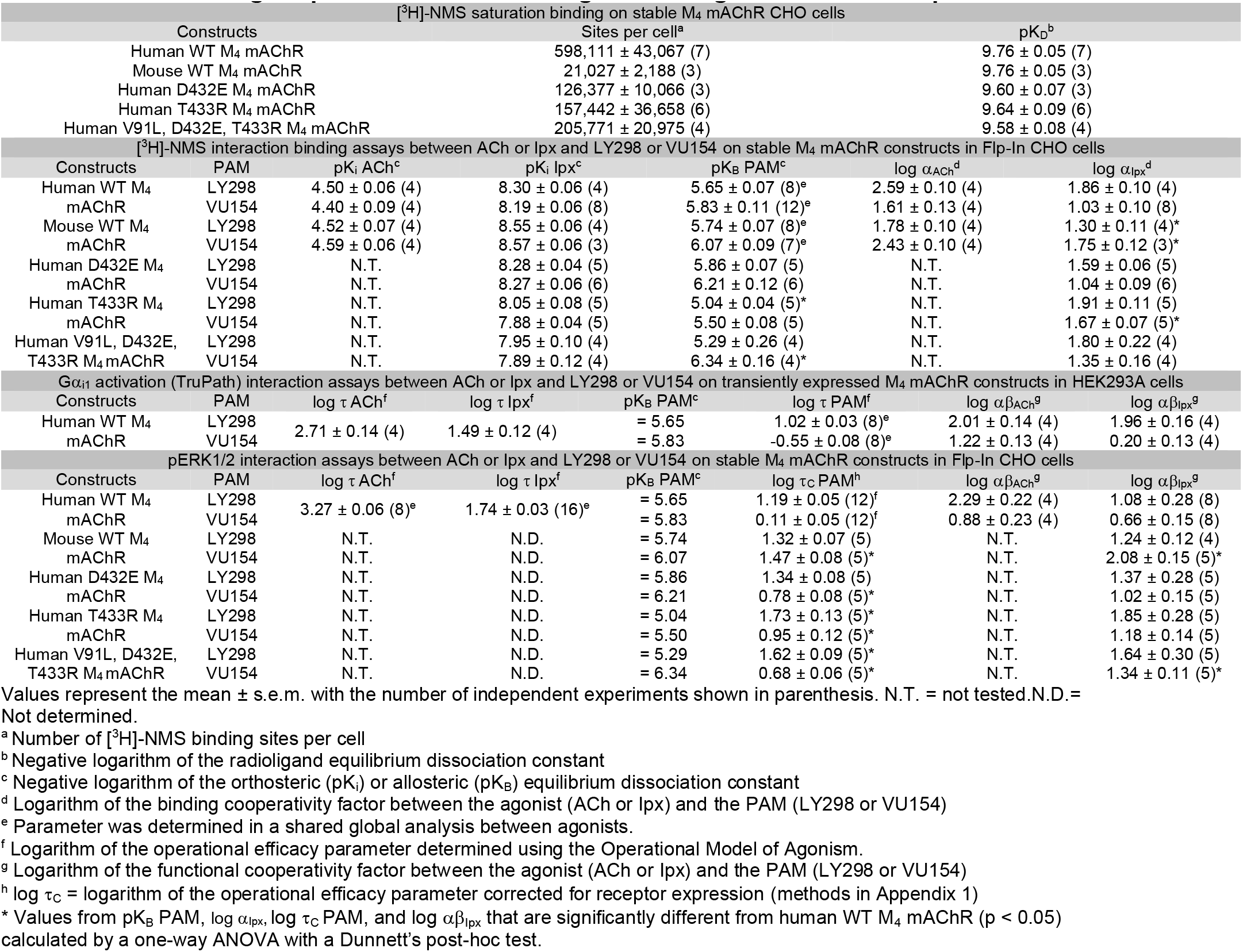
Pharmacological parameters from radioligand binding and functional experiments.

**Table S2.**
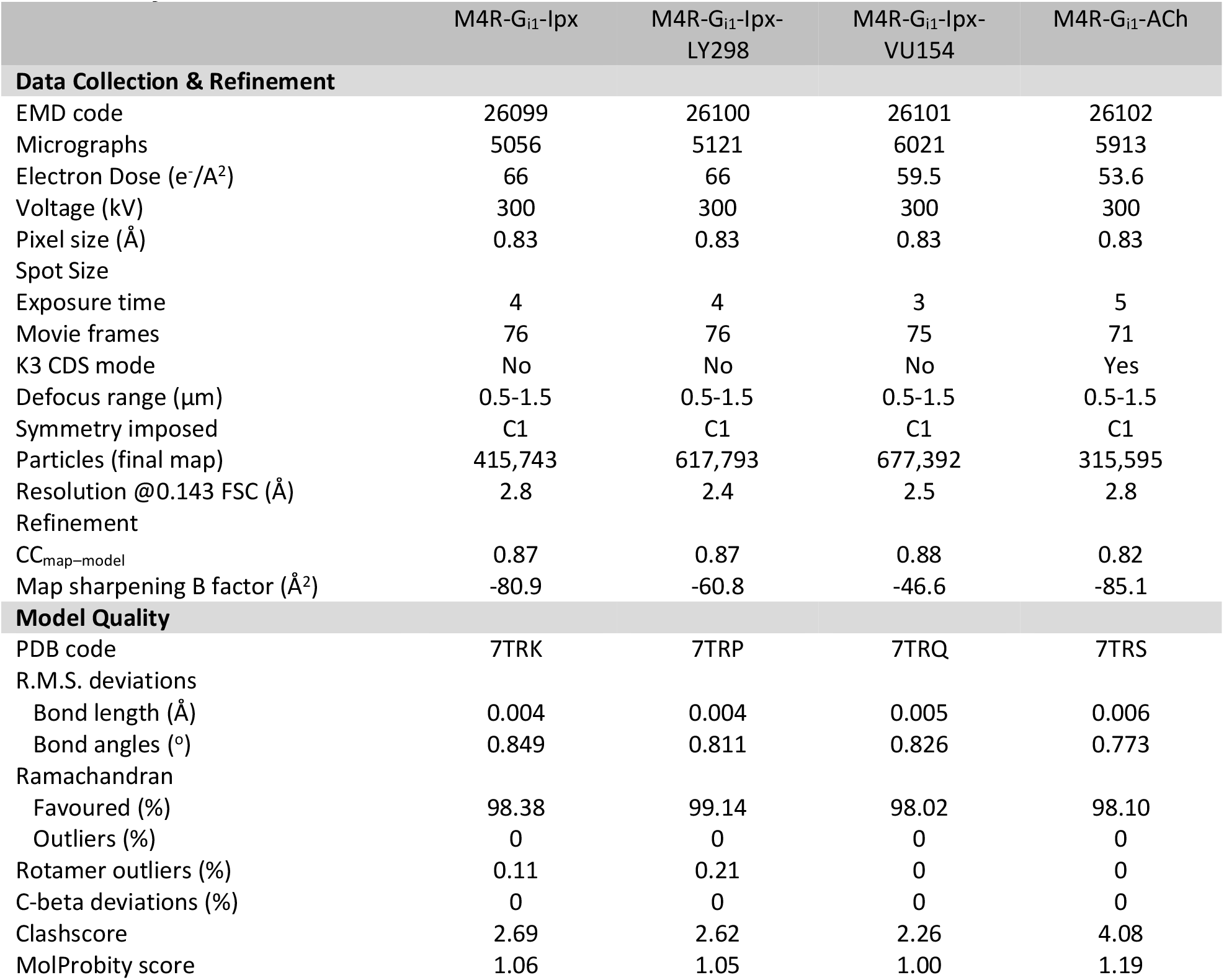
Cryo-EM data collection, refinement, and validation statistics

**Table S3:** GaMD simulations of the M_4_ mAChR

| System | Method |
| --- | --- |
| M4-G<sub>i1</sub>-lpx (cryo-EM structure) | GaMD (3 x 500 ns) |
| M4-G<sub>i1</sub>-lpx-VU154 (cryo-EM structure) | GaMD (3 x 500 ns) |
| M4-G<sub>i1</sub>-lpx-LY298 (cryo-EM structure) | GaMD (3 x 500 ns) |
| M4-G<sub>i1</sub>-ACh (cryo-EM structure) | GaMD (3 x 500 ns) |
| M4-D432E-G<sub>i1</sub>-lpx-VU154 | GaMD (3 x 500 ns) |
| M4-T433R-G<sub>i1</sub>-lpx-VU154 | GaMD (3 x 500 ns) |
| M4-G<sub>i1</sub>-ACh-VU154 | GaMD (3 x 500 ns) |
| M4-G<sub>i1</sub>-ACh-LY298 | GaMD (3 x 500 ns) |
| M4-G<sub>i1</sub>-VU154 | GaMD (3 x 500 ns) |
| M4-G<sub>i1</sub>-LY298 | GaMD (3 x 500 ns) |
| M4-VU154 | GaMD (3 x 1,000 ns) |
| M4-LY298 | GaMD (3 x 1,000 ns) |

**Table S4.**
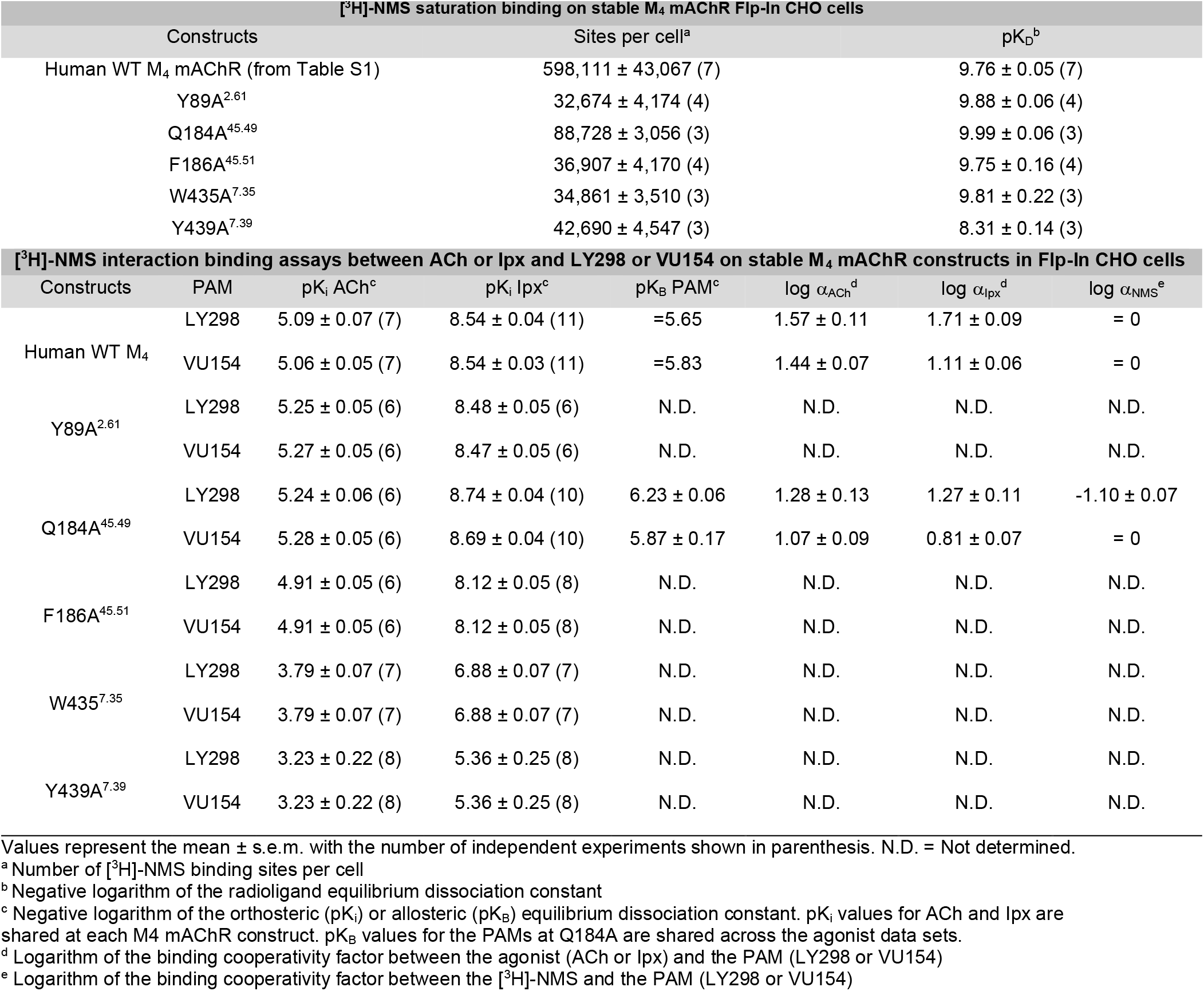
Pharmacological parameters of LY298 and VU154 at key M_4_ mAChR mutants.

---

**Supplementary Video 1. 3D variability analysis of the Ipx-M_4_R-G_i1_ cryo-EM structure**

**Supplementary Video 2. 3D variability analysis of the ACh-M_4_R-G_i1_ cryo-EM structure**

**Supplementary Video 3. 3D variability analysis of the LY298-Ipx-M_4_R-G_i1_ cryo-EM structure**

**Supplementary Video 4. 3D variability analysis of the VU154-Ipx-M_4_R-G_i1_ cryo-EM structure**

**Supplementary Video 5. Movie from one Ipx-M_4_R-G_i1_ GaMD simulation**

**Supplementary Video 6. Movie from one LY298-Ipx-M_4_R-G_i1_ GaMD simulation**

**Supplementary Video 7. Movie from one VU154-Ipx-M_4_R-G_i1_ GaMD simulation**

**Supplementary Video 8. Movie from one ACh-M_4_R-G_i1_ GaMD simulation**

**Supplementary Video 9. Movie from one VU154-Ipx-M_4_R(D432E)-G_i1_ GaMD simulation Supplementary Video 10. Movie from one VU154-Ipx-M4R(T433R)-G_i1_ GaMD simulation**

